# The neuroblast timer gene *nubbin* exhibits functional redundancy with gap genes to regulate segment identity in *Tribolium*

**DOI:** 10.1101/2021.04.08.438913

**Authors:** Olivia RA Tidswell, Matthew A Benton, Michael E Akam

## Abstract

In *Drosophila*, segmentation genes of the gap class form a regulatory network that positions segment boundaries and assigns segment identities. This gene network shows striking parallels with another gene network known as the neuroblast timer series. The neuroblast timer genes *hunchback, Krüppel, nubbin,* and *castor* are expressed in temporal sequence in neural stem cells to regulate the fate of their progeny. These same four genes are expressed in corresponding spatial sequence along the *Drosophila* blastoderm. The first two, *hunchback* and *Krüppel*, are canonical gap genes, but *nubbin* and *castor* have limited or no roles in *Drosophila* segmentation. Whether *nubbin* and *castor* regulate segmentation in insects with the ancestral, sequential mode of segmentation remains largely unexplored.

We have investigated the expression and functions of *nubbin* and *castor* during segment patterning in the sequentially-segmenting beetle *Tribolium*. Using multiplex fluorescent *in situ* hybridisation, we show that *Tc-hunchback*, *Tc-Krüppel*, *Tc-nubbin* and *Tc-castor* are expressed sequentially in the segment addition zone of *Tribolium*, in the same order as they are expressed in *Drosophila* neuroblasts. Furthermore, simultaneous disruption of multiple genes reveals that *Tc-nubbin* regulates segment identity, but does so redundantly with two previously described gap/gap-like genes, *Tc-giant* and *Tc-knirps*. Knockdown of two or more of these genes results in the formation of up to seven pairs of ectopic legs on abdominal segments. We show that this homeotic transformation is caused by loss of abdominal Hox gene expression, likely due to expanded *Tc-Krüppel* expression. Our findings support the theory that the neuroblast timer series was co-opted for use in insect segment patterning, and contribute to our growing understanding of the evolution and function of the gap gene network outside of *Drosophila*.

## Introduction

The gap gene network of *Drosophila* is arguably one of the best characterised gene regulatory networks in developmental biology. Gap genes mediate two central processes in *Drosophila* segmentation – the formation of segment boundaries and the assignment of segment identities - through direct regulation of pair-rule and Hox genes, respectively (reviewed in [1]). Homologs of many *Drosophila* gap genes also regulate segment patterning in other insect species [2–8]. Recent attention has therefore turned to understanding how gap genes interact and function outside of *Drosophila*, in order to better understand the origins and evolution of this important gene network.

In *Drosophila*, the gap genes are thought of as markers for spatial domains, regulated initially by gradients of maternal factors, and then by cross regulation within the gap gene network itself [1]. However, recent work, particularly in the red flour beetle *Tribolium castaneum*, leads to a rather different way of viewing these same genes. In *Tribolium* and other sequentially segmenting insects, segments are added progressively, from anterior to posterior, from a segment addition zone (SAZ) at the posterior of the extending germ band [9]. Gap genes are sequentially activated in the SAZ, so that cells persisting in this region experience a temporal sequence of gap gene expression [10,11] (Fig 1, A1-A2). As each cell exits the SAZ, its gap gene expression is stabilised [11], creating a spatial pattern of gap gene expression along the anterior-posterior axis of the trunk. The gap genes may therefore provide a timer for the maturation of cells with different axial identities from the segment addition zone [2,3,9].

**Fig 1.**
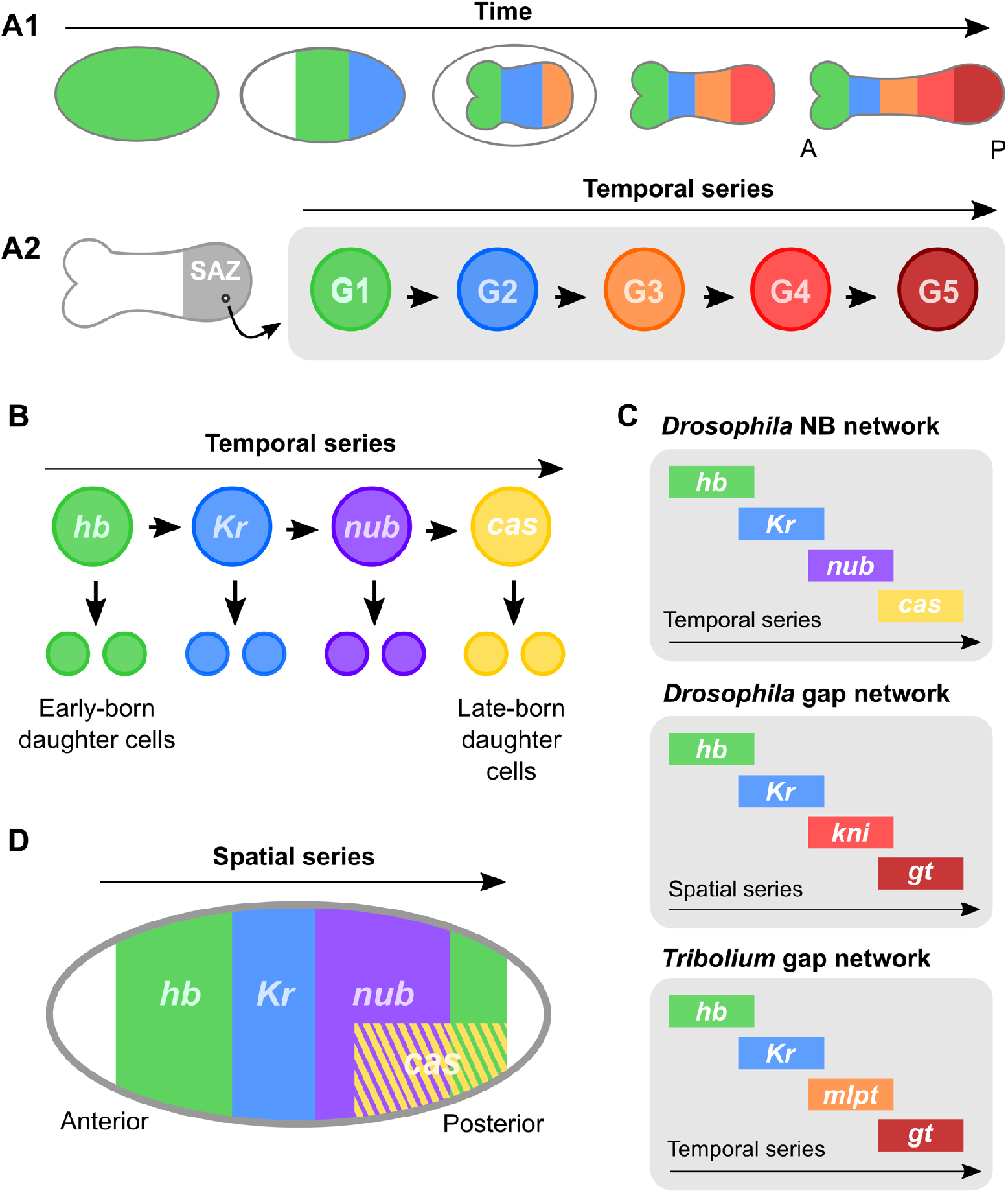
Parallels between the gap gene network and neuroblast timer network in insects. **(A1)** The expression domains of gap genes emerge sequentially from the posterior of the embryo (the SAZ) in sequentially-segmenting insects. **(A2)** Cell lineages persisting in the SAZ therefore express a temporal sequence of gap genes. **(B)** The neuroblast timer genes *hb*, *Kr*, *nub* and *cas* are expressed sequentially in embryonic neuroblasts of *Drosophila*, where they regulate the assignment of daughter cell fates. **(C)** A side-by-side comparison of the series of genes expressed in the *Drosophila* neuroblast (NB) timer network and the canonical *Drosophila* and *Tribolium* gap gene networks. **(D)** The neuroblast timer genes, including *Dm-nub* and *Dm-cas*, are expressed along the anterior-to-posterior (AP) axis of the *Drosophila* blastoderm in roughly the same spatial order as they are expressed temporally in neuroblasts. A= anterior, P = posterior, G1-5 = gap genes 1-5.

This model of the gap gene network has many similarities to the neuroblast timer network that regulates embryonic neural patterning in insects [9,12]. The insect nervous system is produced by neural stem cells known as neuroblasts, each of which gives rise to a range of different cell types in a stereotyped order. In embryonic neuroblasts of *Drosophila*, this order is directed by the sequential expression of the neuroblast timer genes *hunchback* (*hb*), *Krüppel* (*Kr*), *nubbin* (*nub*), *castor* (*cas*) and *grainyhead* (*grh*) (reviewed in [13]) (Fig 1, B). Remarkably, homologues of at least *hb*, *nub* and *cas* are expressed in the same relative order in vertebrate neural stem cells, where they regulate the fate of neurons derived from their progeny [14–18]. This suggests that the roles of these genes in neural development are deeply conserved.

Parallels between the neuroblast timer series and the gap gene network have long been noted [19,20] giving rise to the hypothesis that elements of the neuroblast timer network may have been co-opted from neuroblasts for use in insect axial patterning [20]. The first two genes in the neuroblast timer series, *hb* and *Kr,* are also canonical gap genes in *Drosophila* and *Tribolium* [3,6,10]. However, the next two genes in the neuroblast timer series, *nub* and *cas*, are not canonical gap genes in *Drosophila* [1]. The canonical gap genes acting posterior to *Kr* in *Drosophila* - *Dm-knirps* (*kni*) and *Dm-giant* (*gt*) *–* and in *Tribolium – Tc-gt* and *Tc-mille-pattes -* are not components of the neuroblast timer series (Fig 1, C).

While *nub* and *cas* are not canonical gap genes, they do show some intriguing similarities to gap genes. In *Drosophila*, the *nub* gene has been duplicated to give rise to the closely linked genes *pdm1* (*nub*) and *pdm2* with overlapping expression domains [21]. *Dm-nub*/*Dm-pdm2* and *Dm-cas* are expressed in the *Drosophila* blastoderm during segment patterning, in spatial domains that follow in sequence behind *Dm-hb* and *Dm-Kr* [19] (Fig 1, D). Ectopic expression of *Dm-nub* or *Dm-pdm2* results in gap-like segment deletions [21]; however, neither gene appears to regulate the canonical gap genes [21], and deletion of both genes generates only incompletely penetrant and variable segment fusions [21,22]. *Dm-cas* is not known to have any role in segmentation (for example, see [23]).

Data from sequentially segmenting insects has identified further parallels. A homologue of *nub* is necessary for the correct specification of abdominal segment identity in the bug *Oncopeltus* [24], although not in the cricket *Acheta* [25]. In *Tribolium*, *Tc-nub* and *Tc-cas* are also expressed in the segment addition zone [26], but parental RNA interference has failed to identify any role for *Tc-nub* in segmentation (E. Raymond and A. Peel, personal communication), and functional analyses have not been carried out for *Tc-cas*.

In this paper we examine whether *Tc-nub* and *Tc-cas* form part of a temporal sequence of gene expression during segmentation in *Tribolium*, and we ask whether either regulates segment addition or the assignment of segment identities. Our functional analyses demonstrate a clear role for *Tc-nub* in the assignment of abdominal segment identity. This role is partially redundant with that of other abdominal gap genes, explaining why it has not been identified previously. Our findings strengthen the hypothesis that elements of the gap gene network may have been recruited for a timing role in axial extension from a pre-existing role in neural development.

## Results

### The neuroblast timer genes are expressed sequentially in the SAZ

We first examined whether the genes of the neuroblast timer series are expressed in temporal order in the SAZ of *Tribolium* during segment addition. We used Hybridisation Chain Reaction *in situ* hybridisation (HCR ISH) [27] to examine the expression patterns of *Tc-hb*, *Tc-Kr*, *Tc-nub* and *Tc-cas* in *Tribolium* embryos spanning the stages of segment addition (8-22 hours after egg lay (AEL) at 30 °C). We found that these four genes are expressed sequentially in the SAZ in the same order as they are expressed in neuroblasts (Fig 2, A-I), and that this sequential expression results in their being expressed in spatial order along the anterior-to-posterior (AP) axis of the embryonic trunk (Fig 2, J). *Tc-hb* mRNA is initially expressed broadly across the blastoderm [28] (Fig 2, A), becoming lost from the posterior tip of the embryo as *Tc-Kr* expression emerges [10] (Fig 2, B). *Tc-nub* becomes expressed at the posterior tip of the embryo shortly afterwards, correlating with loss of *Tc-Kr* expression in the same region (Fig 2, C-D). *Tc-cas* becomes expressed in the SAZ midway through germband extension, in a domain overlapping with the posterior of the *Tc-nub* domain (Fig 2, E-F). Finally, a second domain of *Tc-hb* becomes expressed in the posterior SAZ and remains expressed in the SAZ until the end of segment addition (Fig 2, E-H). Each gene is also expressed in the neurectoderm and neuroblasts in differentiating segments, as well as in the tissue at the extreme posterior of the embryo (the presumptive hindgut epithelium [29]) after segmentation is complete (S1 Fig).

**Fig 2.**
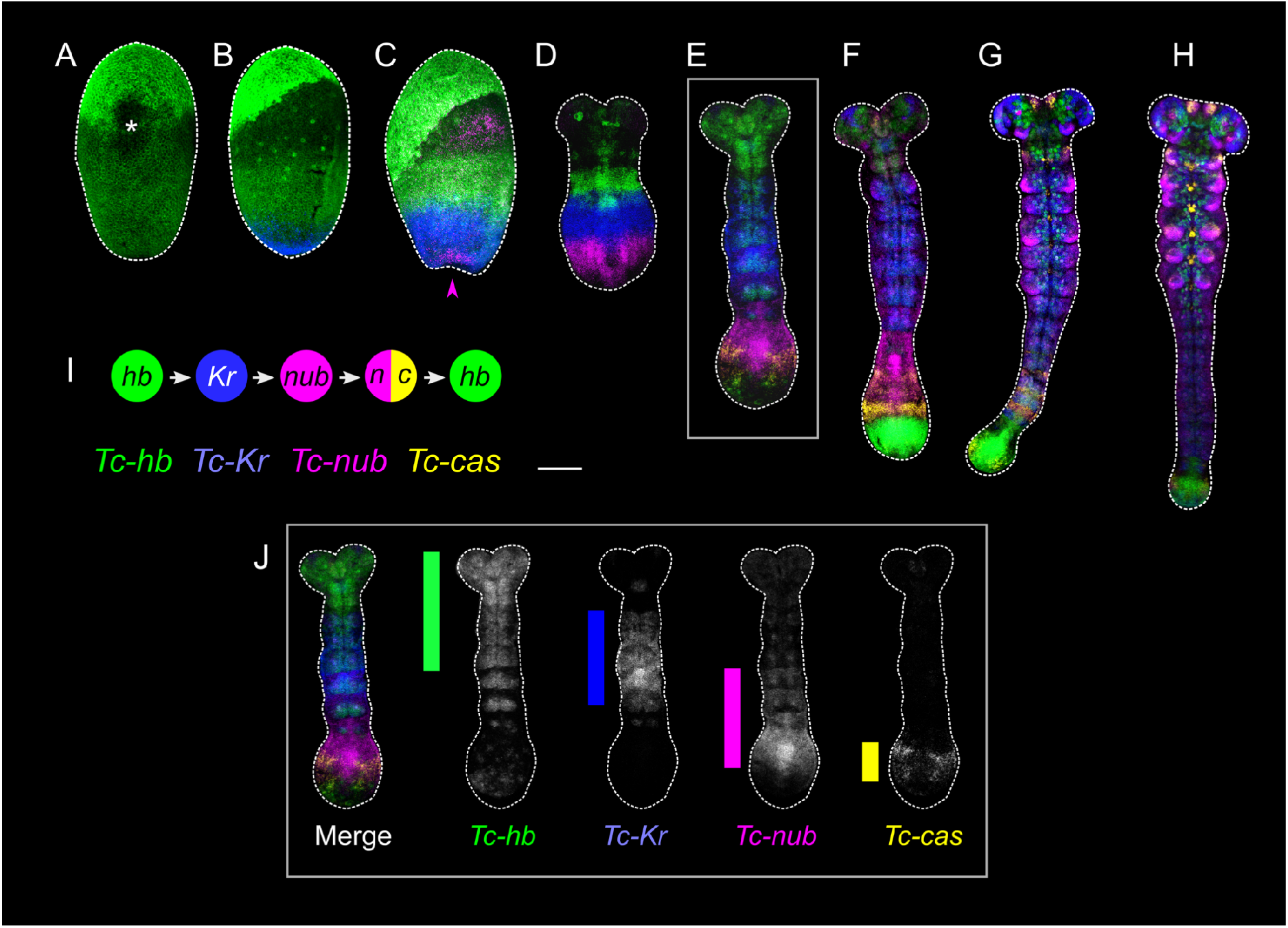
Expression of the neuroblast timer genes *Tc-hb*, *Tc-Kr*, *Tc-nub* and *Tc-cas* during segment addition in *Tribolium*. **(A-H)** HCR ISH was used to visualise the expression of *Tc-hb*, *Tc-Kr*, *Tc-nub* and *Tc-cas* in embryos spanning the course of segment addition, from the differentiated blastoderm stage (A) to the end of segment addition (H). The asterisk in A highlights damage to the embryo. The magenta arrow in C indicates the newly emerging domain of *Tc-nub* expression in the posterior pit. The expression of *Tc*-*hb*, *Tc*-*Kr* and *Tc-nub* outside of the SAZ in later embryos (E-H) can be attributed at least partly to roles in other embryological processes such as mesodermal and neural patterning. **(I)** A graphical summary of the gene expression states experienced by cells in the SAZ, inferred from the expression dynamics in A-H. **(J)** Greyscale images showing the expression of *Tc-hb*, *Tc-Kr*, *Tc-nub* and *Tc-cas* in embryo E. Coloured bars are used to highlight the extent of their “gap” expression domains along the AP axis (excluding segments where expression is restricted to the mesoderm or neuroectoderm). All images are maximum projections of confocal z-stacks through the top half of the embryo (A-C) or through the entire embryo (D-H and J). Dissected germbands (D-H and J) are flat-mounted. In all panels, anterior is to the top. Ventral is along the vertical midline of each image except in B and C, where it is towards the right. Scale bar is 100 μM.

### Expression of *Tc-nub* and *Tc-cas* in relation to segment patterning

To characterise the expression dynamics of *Tc-nub* and *Tc-cas* in more detail, we next examined the expression of both genes against expression of the segment polarity gene *Tc-wingless* (*Tc-wg*) [30] in embryos spanning the course of segment addition. The number of *Tc-wg* stripes in each embryo is used as an indicator of the progression of segment addition, and therefore as a proxy for developmental age.

The earliest expression of *Tc-nub* is at the late blastoderm stage in two patches overlying the ocular *Tc-wg* stripes (Fig 3, A1). It is first expressed at the posterior pole shortly afterwards (Fig 3, B1). This posterior domain subsequently expands to encompass the posterior third of the SAZ as the embryo condenses to form a germband (Fig 3, C1). The anterior border of this broad, gap-like domain abuts the *Tc-wg* stripe at the posterior of parasegment 3 (PS3), which we refer to as wg3 (Fig 3, D1). Expression is weakest in the anterior of this domain, and strongest in the posterior SAZ. After the formation of wg6, *Tc-nub* expression begins to fade in the posterior SAZ, and the posterior boundary shifts anteriorly to overlap with wg12 (Fig 3, F1-J1). *Tc-nub* is therefore expressed in the SAZ during the patterning of PS4-PS12 (posterior compartment of T1 to anterior compartment of A7, inclusive). This overlaps extensively with the expression of the gap genes *Tc-mlpt*, *Tc-gt* and the gap-like gene *Tc-kni* (Fig 3, L1).

**Fig 3.**
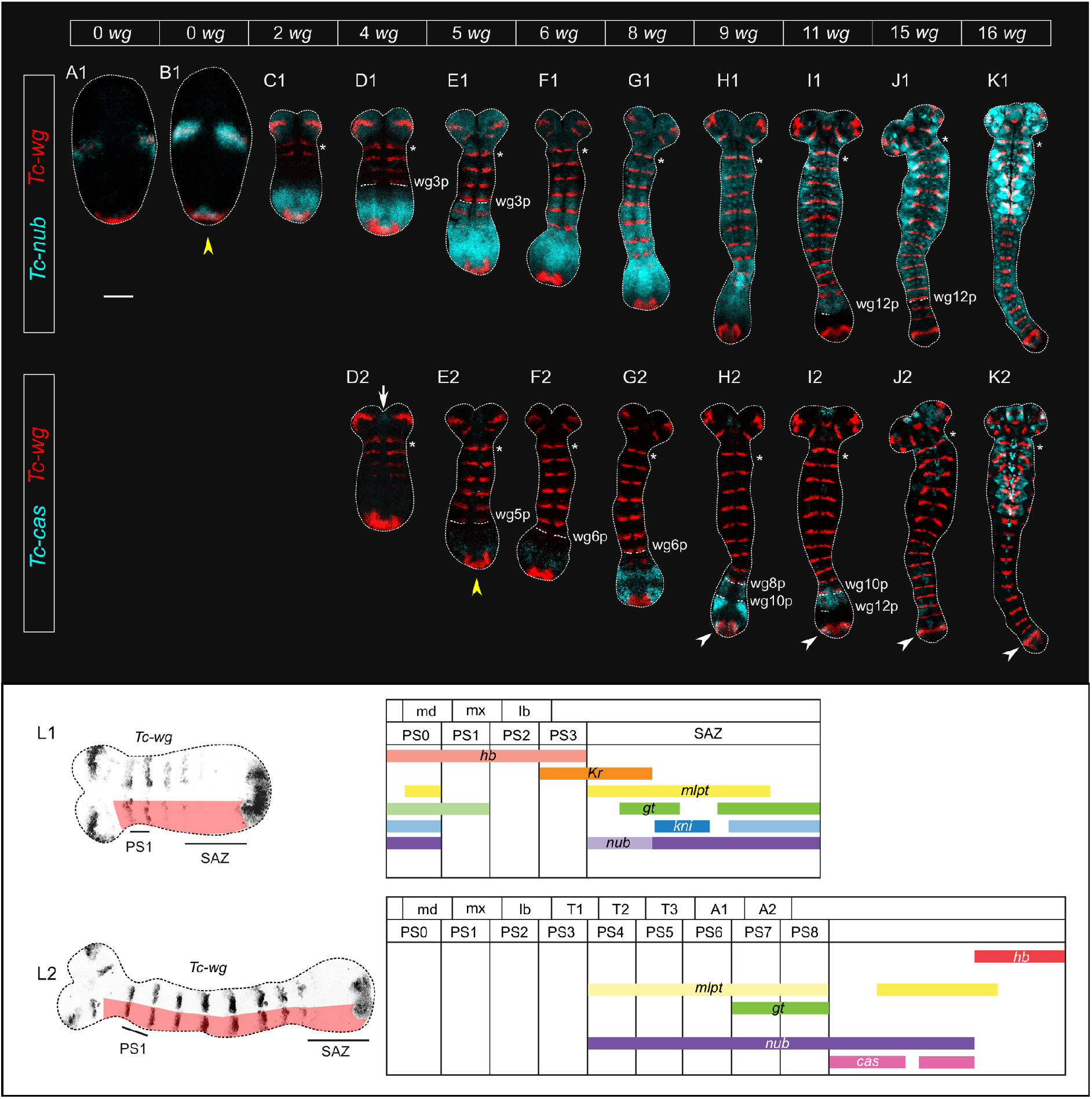
Expression of *Tc-nub* and *Tc-cas* in *Tribolium* embryos during segment addition, using *Tc-wg* as a segmental marker. **(A-K)** HCR ISH was used to examine the expression of *Tc-nub* (A1-K1) and *Tc-cas* (D2-K2) over the course of segment addition. Images in the same column come from the same embryo. Asterisks mark the first *Tc-wg* stripe to form in the trunk, which we refer to as wg0 in reference to its location within the posterior of parasegment (PS) 0. wg3-12p *=* the posterior boundary of *Tc-wg* stripes 3-12. Yellow arrowheads mark the onset of expression of *Tc-nub* and *Tc-cas* in the segment addition zone. The white arrow in D2 indicates faint *Tc-cas* expression in the developing labrum, and the white arrowheads in H2-K2 indicate the domain of *Tc-cas* expression that overlaps with the terminal domain of *Tc-Wg* expression. (**L1-L2**) Diagrams of *Tc-nub* and *Tc-cas* expression relative to the expression of other *Tribolium* gap genes (based on published descriptions [2,3,27,34,41,53]) in embryos with 4 (L1) or 9 (L2) trunk *Tc-wg* stripes. These abstractions span from a short distance anteriorly of wg0 (*i.e*., within PS0) to the anterior boundary of the terminal domain of *Tc-wg* (as indicated with red shading on the reference embryos to the left). md = mandibular segment, mx = maxillary segment, lb = labial segment, T1-T3 = thoracic segments 1-3, A1-A2 = abdominal segments 1-2. All images are maximum projections of confocal z-stacks through half of the embryo (A1 and B1) or through the entire embryo (C1-L2). Dissected germbands (C1-L2) are flat-mounted. In (A1-K2) anterior is to the top, while in (L1-L2) anterior is to the left. Ventral is along the vertical (A1-K2) or horizontal (L1-L2) midline of each image. Scale bar is 100 μM.

*Tc-cas* expression is not detectable in the embryo until after the germband has formed. It is initially expressed weakly in the primordium of the labrum (Fig 3, D2), before becoming weakly expressed in the posterior SAZ after the formation of wg5 (Fig 3, E2-F2). The anterior border of this expression domain abuts the posterior boundary of wg6 (Fig 3, F2-G2). Within the SAZ, the expression of *Tc-cas* is modulated in a pair-rule pattern, with the strongest domains of expression overlapping approximately the primordia for PS9 and PS11 (Fig 3, H2). Expression of *Tc-cas* subsequently fades in the posterior SAZ (Fig 3, H2) and the posterior boundary shifts anteriorly to overlap with wg12 (Fig 3, I2). This means that *Tc-cas* is expressed in the SAZ during the patterning of PS7-12 (posterior compartment of A3 to anterior compartment of A7), overlapping with expression of *Tc-nub* and *Tc-mlpt* (Fig 3, L2). Outside of the SAZ, in maturing segments, *Tc-cas* expression fades and is lost (Figure 3, G2-J2). There is an additional domain of *Tc-cas* that overlaps with the posterior terminal domain of *Tc-wg* (Figure 3, H2-K2).

*Tc-nub* and *Tc-cas* are both also expressed in the developing neuroectoderm and neuroblasts [26] (Figure 3, G1-K1 and J2-K2).

### *Tc-nub*, but not *Tc-cas,* influences segment identity

We next aimed to determine whether *Tc-nub* or *Tc-cas* have a role in axial patterning in *Tribolium*. To do this, we knocked down the expression of each gene by parental and embryonic RNA interference (pRNAi and eRNAi, respectively).

We found that pRNAi and eRNAi against *Tc-nub* (2 μg/μL dsRNA) resulted in a subtle abdominal segment transformation in a small percentage of embryos. Specifically, 1.7% (2/120, pRNAi) and 7.4% (11/148, eRNAi) of embryos displayed a ‘nub’ (an ectopic protrusion of cuticle, lacking joints or claws) on either side of segment A1 (Figure 4, B; S1 and S2 Tables). Similar nubs form following pRNAi against *Tc-abdominal-A* (*Tc-abd-A,* also known as *Tc-Abdominal* [31]), and have been interpreted as homeotic transformations of the posterior compartment of an abdominal segment to the posterior compartment of segment T3 [32]. This would make each nub developmentally akin to the posterior compartment of a thoracic leg. We examined the expression of *Tc-abd-A* in our RNAi embryos, and observed that a small percentage of embryos showed a loss or reduction of expression of *Tc-abd-A* in the anterior of PS7, which gives rise to A1p (S2 Fig).

**Fig 4.**
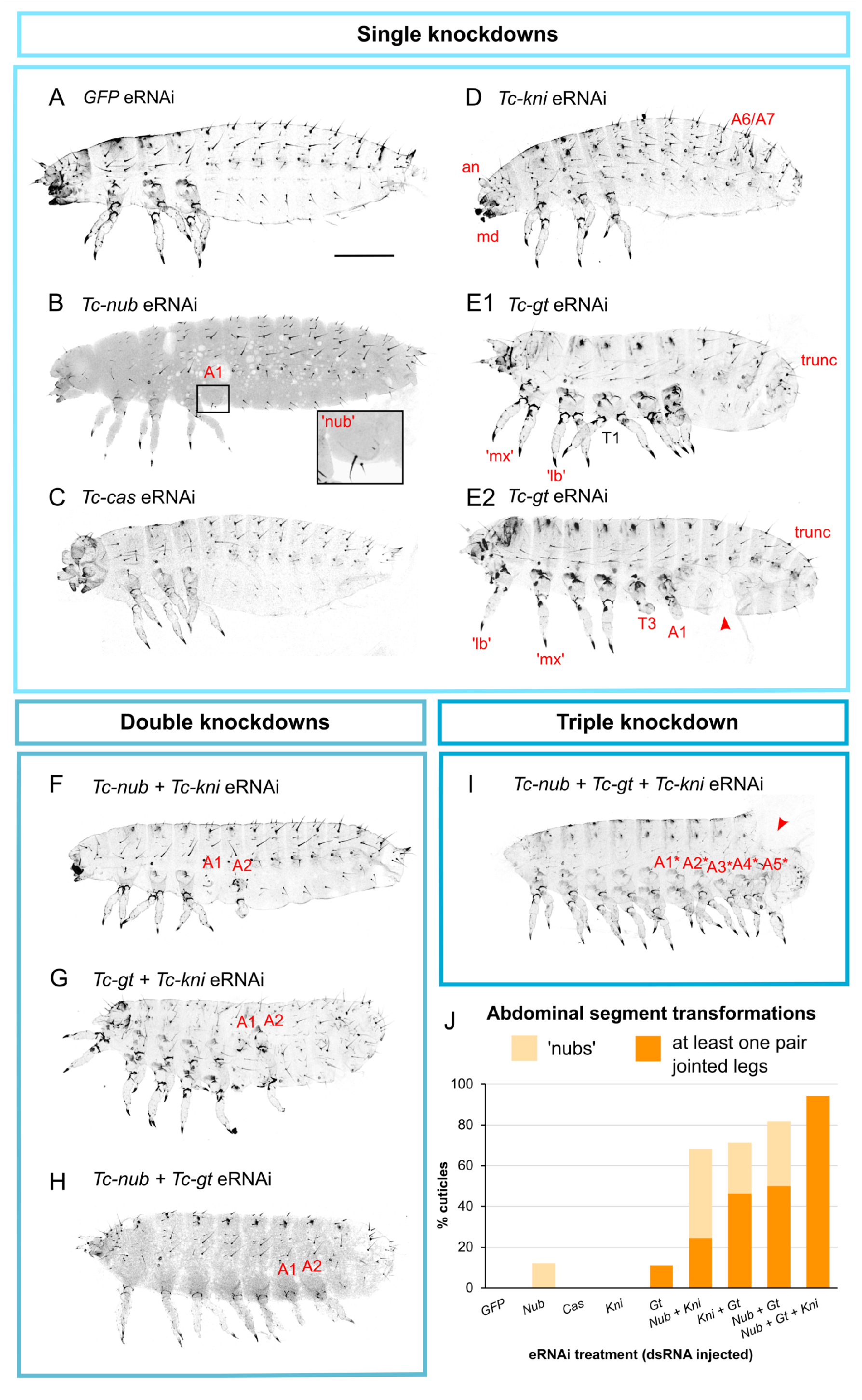
*Tc-nub* acts redundantly with *Tc-gt* and *Tc-kni* to regulate abdominal segment identity. **(A)** Control embryos injected with *GFP* dsRNA (2 μg/μL) displayed wild type abdominal segment morphology. **(B)** A small percentage of embryos injected with *Tc-nub* dsRNA ( 2 μg/μL) formed cuticular protrusions (nubs, magnified in the inset) on the first abdominal segment (A1). **(C)** Embryos injected with *Tc-cas* dsRNA (2 μg/μL) showed no consistent defects in cuticular morphology. This specific embryo displays head defects that were common in all treatments, probably resulting from the injections at the anterior pole of the embryo. **(D)** Embryos injected with *Tc-kni* dsRNA (2 μg/μL) frequently lacked antennal (an) and/or mandibular (md) segments, and displayed disrupted segment patterning in the posterior abdomen. **(E1-E2)** Embryos injected with *Tc-gt* dsRNA (2 μg/μL) formed thoracic legs in the place of maxilla (mx) and labium (lb), and displayed posterior truncation of the abdomen (trunc). We also observed that a small percentage of injected embryos developed ectopic legs on segment A1 (E2). **(F-H)** Embryos injected with any two of *Tc-nub*, *Tc-gt* and *Tc-kni* dsRNAs (1 μg/μL each) frequently formed cuticular protrusions (nubs) and/or ectopic legs (with joints and/or claws) on segments A1 and/or A2. **(I)** Embryos injected with *Tc-nub*, *Tc-gt* and *Tc-kni* dsRNA (1 μg/μL each) formed ectopic legs on the majority of abdominal segments. The red arrow indicates damage to the cuticle sustained during dissection from the eggshell. The asterisks in G indicate that these segment assignments are estimates, as we cannot be entirely certain how many head segments are deleted. Note that the cuticles in F-I also display head defects consistent with the repression of *Tc-gt* and/or *Tc-kni* expression. All images are maximum projections of confocal z-stacks through cuticle preparations. Anterior is to the left, and dorsal is to the top. **(J)** A bar graph summarizing the frequency of ‘weak’ transformations (displaying only nubs) compared to ‘strong’ transformations (displaying at least one pair of ectopic, jointed legs) on abdominal segments following various eRNAi treatments. Numbers of cuticles examined and phenotypes observed for each treatment are provided in S2 Table. Scale bar is 200 μM.

Neither pRNAi nor eRNAi against *Tc-cas* had any consistent effects on cuticular morphology or on segment patterning in embryos (Figure 4, C and S1 Table; N=116 and 89 embryos examined, respectively).

Although the vast majority of larvae that developed following *Tc-nub* or *Tc-cas* pRNAi had wild type external morphology, they displayed a severely reduced hatching rate compared to injection controls (2-4% of *Tc-nub* or *Tc-cas* knockdowns hatched successfully, compared to 82% of injection controls) (S3 Fig). Reduced hatching rate was also observed at a lower dsRNA concentration (1 μg/μL), when the nub phenotype was no longer observed (S1 Table). The failure of otherwise ‘normal’ larvae to hatch could be a result of defects in the nervous system. Both *Tc-nub* and *Tc-cas* are expressed in neuroblasts in *Tribolium* [26], and mutation of *Dm-cas* has been shown to prevent hatching of *Drosophila* embryos with otherwise normal cuticles, presumably because of disruption to the nervous system [23].

We also found that pRNAi, but not eRNAi, knockdown of *Tc-nub* or *Tc-cas* significantly reduced the proportion of eggs that developed to the stage of cuticle formation compared to injection controls (S3 Fig). *Tc-nub* and *Tc-cas* are both expressed in the ovarioles of adult female *Tribolium* (S4 Fig). We therefore suspect that both genes have roles in oogenesis or early embryogenesis in *Tribolium*. *Dm-cas* is known to be required for the proper formation of follicular cells [33] but *Dm-nub* does not seem to be expressed in ovaries [34].

Together, our data show that *Tc-nub* and *Tc-cas* are likely involved in oogenesis and neurogenesis, and that *Tc-nub* affects specification of segment identity.

### *Tc-nub* acts redundantly with *Tc-gt* and *Tc-kni* to regulate abdominal segment identity

The spatially restricted and weakly penetrant homeotic phenotype observed after *Tc-nub* RNAi contrasts with the expression of this gene across the majority of the abdomen. We therefore asked whether *Tc-nub* functions redundantly with other gap genes. Two promising candidate genes for redundant function are *Tc-giant* (*Tc-gt*) and *Tc-knirps* (*Tc-kni*), both of which are transiently co-expressed with *Tc-nub* in the SAZ (Fig 3, L1; S5 Fig). *Tc-gt* is considered a gap gene in *Tribolium*, as its knockdown affects thoracic segment identity and abdominal segment formation [2]. In contrast, *Tc-kni* is not considered to be a gap gene, as its knockdown results in the deletion of only one segment boundary in the head, with no effects on segment identity [35,36].

To determine whether *Tc-nub* acts redundantly with *Tc-gt* and/or *Tc-kni* to regulate abdominal segment patterning, we performed single, double and triple knockdowns of these genes. We used eRNAi to avoid the aforementioned negative effects of parental *Tc-nub* knockdown on oogenesis. Single knockdowns of *Tc-kni* and *Tc-gt* produced phenotypes largely consistent with previous reports. Specifically, knockdown of *Tc-kni* resulted in deletions of the antennae and mandibles, and segment patterning defects (most commonly partial fusions) in the region of the abdomen spanning A5-A8; while knockdown of *Tc-gt* resulted in transformation of the maxillary and labial segments to thoracic segments, and axial truncation [2,35,36] (Figure 4, D-E1). The notable exception to previous reports was that 4/36 (11% of) cuticles formed after *Tc-gt* eRNAi also displayed disrupted leg formation on segment T3 and ectopic legs, similarly disrupted, on segment A1 (Figure 4, E2 and J; S2 Table). This difference may be due to eRNAi causing stronger knockdown phenotypes than pRNAi, as has been observed previously [37].

While knockdown of *Tc-nub* or *Tc-gt* alone resulted in only a low frequency of homeotic transformations restricted to A1, and knockdown of *Tc-kni* had no effect on abdominal segment identity, we found that combinatorial knockdown of two or more of these genes generated more severe phenotypes at a higher frequency (Fig 4; S2 Table). After eRNAi against both *Tc-nub* and *Tc-kni*, 18/41 (43% of) cuticles displayed nubs on segments A1 and/or A2, and an additional 10/41 (24% of) cuticles displayed more complete legs (*i.e.* with joints and sometimes claws) on segment A2 (Figure 4, F and J; S2 Table). eRNAi against both *Tc-gt* and *Tc-kni* resulted in 7/28 (25% of) cuticles developing nubs on segments A1 and/or A2, with an additional 13/28 (46% of) cuticles displaying more complete legs on A1 and/or A2 (Figure 4, G and J; S2 Table). Likewise, eRNAi against both *Tc-nub* and *Tc-gt* produced phenotypes that are more severe than can be accounted for additively –12/38 (32% of) cuticles displayed nubs on segments A1 and/or A2, while an additional 19/38 (50%) showed at least one abdominal segment with more complete legs on A1 and/or A2 (Figure 4, H and J; S2 Table).

Finally, knocking down all three genes together produced the most severe and penetrant phenotypes: 33/35 (94% of) cuticles developing from embryos injected with all three dsRNAs formed jointed, clawed legs on at least one abdominal segment (Figure 4, I and J; S2 Table). These cuticles had an average of 4 extra pairs of partial or complete legs (not including the maxillary and labial legs induced by *Tc-gt* knockdown), and a maximum of seven extra pairs (S2 Table), indicating homeotic transformation of up to seven abdominal segments.

In addition to homeotic transformations, *Tc-gt* knockdowns result in truncations of the posterior abdomen with a very high penetrance [2]. In our *Tc-gt* eRNAi experiment, these truncations were observed in 33/35 (94% of) cuticles. Knocking down *Tc-kni* and/or *Tc-nub* in addition to *Tc-gt* did not increase the penetrance or severity of these embryonic truncations (Table 1). Moreover, simultaneous knockdown of *Tc-nub* and *Tc-kni* resulted in a very low frequency of truncations (3/43, 7%), not much higher than what is observed in *GFP* dsRNA controls (1/42, 2%). These data suggest that the truncations observed in *Tc-nub + Tc-gt* or *Tc-nub + Tc-gt + Tc-kni* knockdowns result primarily from loss of *Tc-gt,* and that neither *Tc-nub* nor *Tc-kni* play any substantial role in segment addition. One caveat to this conclusion is that a higher proportion of triple knockdown embryos died before forming cuticle, as compared with double knockdowns (S2 Table), and we observed that many triple knockdown embryos displayed severely disrupted patterning of *Tc-wg* stripes (Fig 6). Therefore, it may be that functional reduction/removal of all three genes has severe effects on the process of segment addition, or other aspects of embryonic growth, that are masked by embryonic death.

**Table 1.** Knockdown of *Tc-nub* and *Tc-kni* does not enhance the severity or penetrance of segment truncations observed after *Tc-gt* knockdown. Single knockdowns were carried out using 2 μg/μL of dsRNA, while all doubles and triple knockdowns used the component dsRNAs mixed to a final concentration of 1 μg/μL each. N cuticles = total number of cuticles examined; # (%) truncated = number (and percentage, in brackets) of cuticles with at least one posterior segment deleted; Avg (Max) # deleted segments = the average (and maximum, in brackets) number of posterior segments deleted over all cuticles examined.

| Treatment (dsRNA injected) |  | N cuticles | # (%) truncated | Avg (Max)<br># deleted segments |
| --- | --- | --- | --- | --- |
| **Singles** | *GFP* | 42 | 2 (1) | 0 (1) |
|  | *Tc-gt* | 34 | 32 (**94**) | **4 (7)** |
| **Doubles** | *Tc-nub* + *Tc-kni* | 43 | 3 (7) | 0 (1) |
|  | *Tc-nub* + *Tc-gt* | 22 | 21 (**95**) | **3 (7)** |
| **Triple** | *Tc-nub* + *Tc-gt* + *Tc-kni* | 19 | 18 (**95**) | **3 (7)** |

*Tc-gt*, *Tc-nub* and *Tc-kni* are also co-expressed during head patterning. However, knocking down two or all three of these genes in parallel did not increase the penetrance or severity of head phenotypes – rather, knockdown effects were additive (S6 Fig), as might be expected if all three genes act independently.

### *Tc-nub*, *Tc-gt* and *Tc-kni* affect segment identity via Hox gene regulation

Development of partial or complete legs on abdominal segments has also been observed in double knockdowns of two abdominal Hox genes, *Tc-abd-A* and *Tc-Ultrabithorax* (*Tc-Ubx,* also known as *Tc-Ultrathorax* [38]) [32]. To determine whether these Hox genes are misexpressed after eRNAi against *Tc-nub*, *Tc-kni* and *Tc-gt*, we performed HCR ISH in embryos midway through segment addition (16-17h AEL). This time point is shortly after the period during which *Tc-nub*, *Tc-gt* and *Tc-kni* are co-expressed, and should, therefore, reveal the immediate effects of knockdown on Hox gene expression.

In wild type embryos, expression of *Tc-Ubx* and *Tc-abdA* is detectable in the SAZ after the formation of the wg2 and wg4, respectively [38,39]. Accordingly, we observed strong expression of both genes in the SAZ of control embryos (injected with GFP dsRNA) just after the formation of wg6 (Fig 5, A1-A4). In contrast, similarly staged embryos injected with *Tc-nub*, *Tc-kni* and *Tc-gt* dsRNA did not express *Tc-Ubx* or *Tc-abdA* (Fig 5, B1-C4). This dramatic loss of Hox gene expression is consistent with the dramatic abdominal phenotypes observed in the cuticles of triple knockdown embryos. Note that the antennal and mandibular *Tc-Wg* stripes (wg0 and wg1) were deleted or highly disorganised in triple knockdowns, consistent with the head cuticle phenotypes observed in Fig S6.

**Fig 5.**
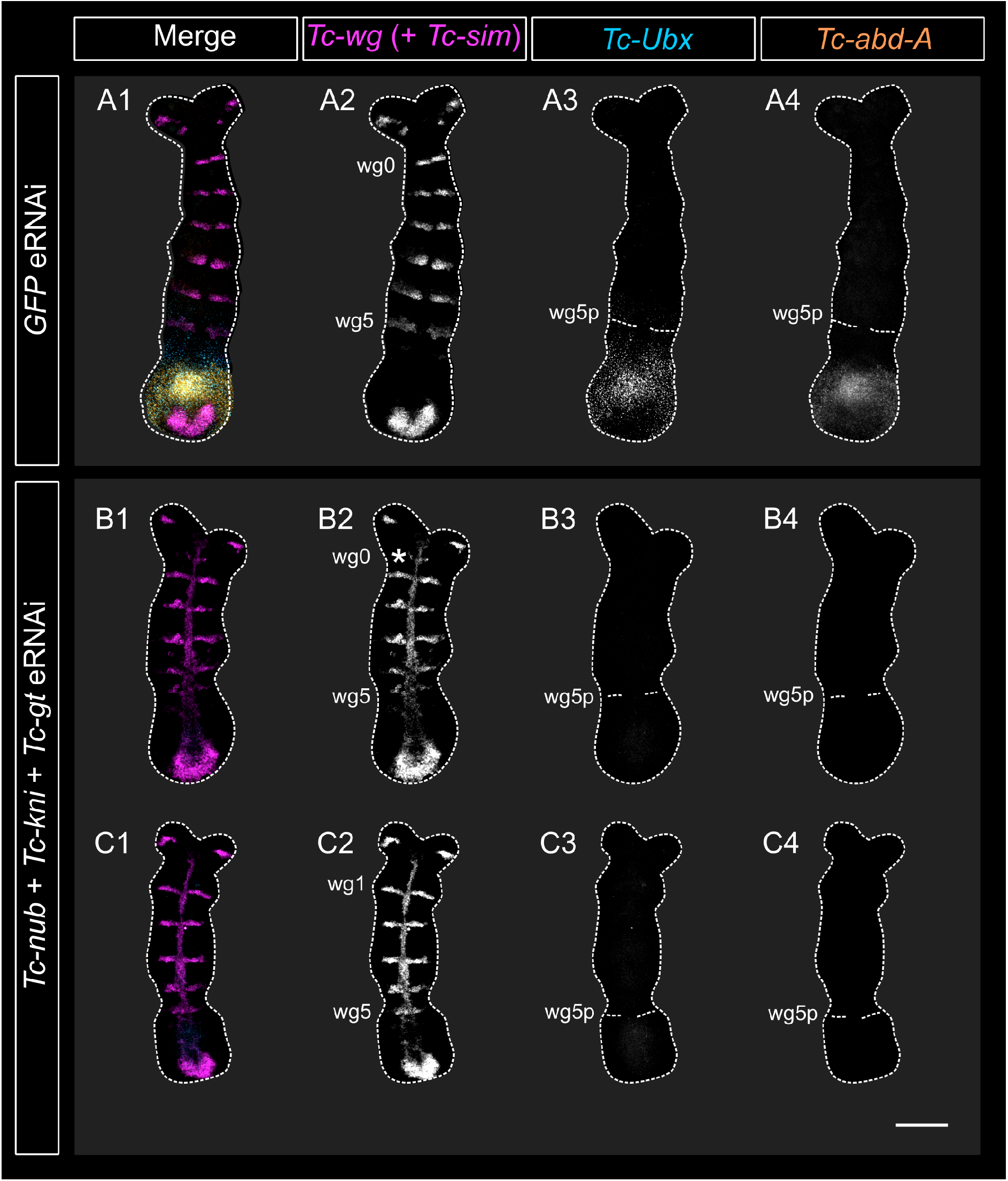
(overleaf). Triple knockdown of *Tc-nub*, *Tc-gt* and *Tc-kni* expression eliminated *Tc-Ubx* and *Tc-abdA* expression in the SAZ. **(A1-A4)** Embryos injected with *GFP* dsRNA (2 μg/μL) expressed *Tc-Ubx* and *Tc-abdA* in the SAZ. **(B1-C4)** At similar stages of segment addition, embryos injected with *Tc-nub, Tc-kni* and *Tc-gt* dsRNA (1 μg/μL each) did not express *Tc-Ubx* or *Tc-abdA* in the SAZ. An asterisk marks what we presume to be the deteriorating mandibular *Tc-Wg* stripe (wg0) in B2, which we believe is deleted entirely in the embryo shown in C1-C4. Note that triple eRNAi embryos were also stained for the expression of the midline marker *Tc-single-minded* in this particular experiment, in the same channel as *Tc-wg*. All embryos were imaged using the same laser settings and brightness/contrast values were adjusted identically for all images. All images are maximum projections of confocal z-stacks through dissected, flat mounted embryos. In all panels, anterior is to the top and ventral is along the vertical midline. wg0-5 *= Tc-wg* stripes 0-5; wg0-5p = posterior boundary of *Tc-wg* stripes 0-5. Scale bar is 100 μM.

**Fig 6.**
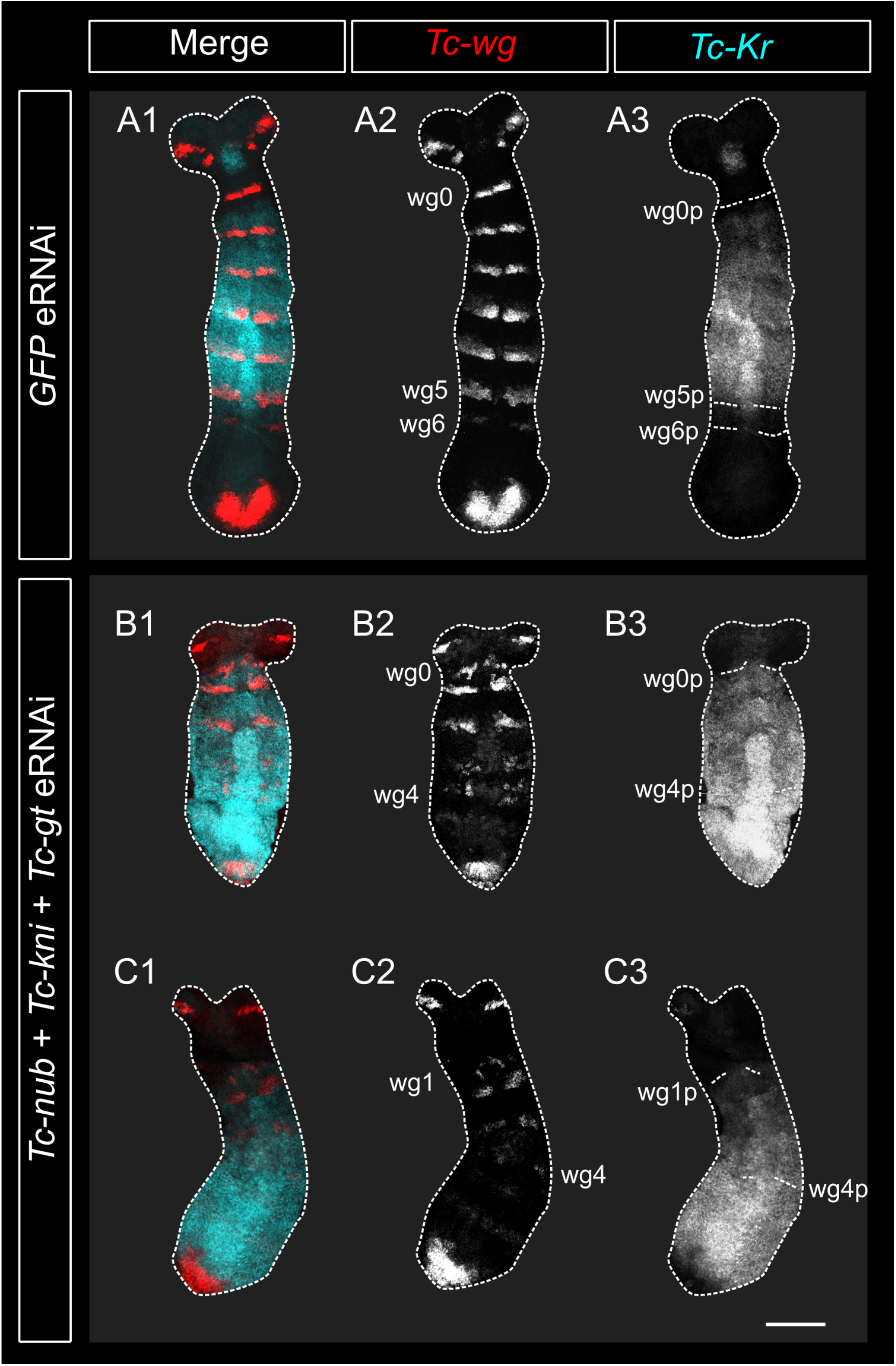
(overleaf). Expression of *Tc-Kr* is expanded posteriorly after knocking down two or more of *Tc-nub*, *Tc-gt* and/or *Tc-kni*. **(A1-A3)** In embryos injected with *GFP* dsRNA (2 μg/μL), *Tc-Kr* expression retracted from the SAZ to cover the presumptive thoracic segments. **(B1-C3)** In embryos injected with *Tc-nub, Tc-kni* and *Tc-gt* dsRNA (1 μg/μL each), *Tc-Kr* failed to retract from the SAZ. Note that the segmental expression of *Tc-wg* was extensively disrupted in the triple knockdown embryos displayed in this figure. We have conservatively estimated that the mandibular stripe (wg0) is intact in B1-B3, but deleted in C1-C3, due to the spacing of stripes relative to the ocular *Tc-wg* stripes in the head. All embryos were imaged using the same laser settings and brightness/contrast values were adjusted identically for all images. All images are maximum projections of confocal z-stacks through dissected, flat mounted embryos. In all panels, anterior is to the top and ventral is along the vertical midline. wg0-6 *= Tc-wg* stripes 0-6; wg0-6p = posterior boundary of *Tc-wg* stripes 0-6. Scale bar is 100 μM.

### *Tc-nub*, *Tc-gt* and *Tc-kni* regulate the expression of *Tc-Kr*, but not *Tc-hb*

We hypothesised that the repression of abdominal Hox genes observed in triple knockdowns might result from the misregulation and expansion of other gap genes. The anterior borders of *Dm-Ubx* and *Dm-abd-A* in *Drosophila* are set primarily via direct repression by *Dm-hb* and *Dm-Kr*, respectively [40,41]. Therefore, we used HCR ISH to examine the expression of both *Tc-hb* and *Tc-Kr* in embryos fixed at 16-17h AEL following eRNAi against *Tc-nub*, *Tc-gt* and *Tc-kni*.

We observed alterations in the pattern of *Tc-Kr*, but not *Tc-hb*, expression in embryos after simultaneous knockdown of *Tc-nub*, *Tc-gt* and *Tc-kni* (Fig 6; S7 Fig). In wild type embryos, *Tc-Kr* is expressed throughout the SAZ at the blastoderm stage, but becomes cleared from the posterior half of the SAZ during early germband formation [35]. This means that the SAZ is largely cleared of *Tc-Kr* expression by the time that the second trunk *Tc-wg* stripe (wg1) is formed [42]. In contrast, triple knockdown embryos with as many as four *Tc-wg* stripes showed little or no clearing of *Tc-Kr* expression in the SAZ (Fig 6, B1-C3). This means that in triple knockdown embryos, *Tc-Kr*, but not *Tc-hb*, is ectopically expressed in the SAZ.

Together, these data suggest that *Tc-nub*, *Tc-Kr* and *Tc-kni* redundantly repress *Tc-Kr* expression, and that in their absence, *Tc-Kr* expression expands into the abdominal primordia. We propose that this expansion leads to the repression of abdominal Hox genes, and subsequently to abdominal segment transformations.

### *Tc-nub* and *Tc-cas* play redundant roles in limb, but not segment, patterning

In addition to double and triple knockdowns of *Tc-nub* with *Tc-gt* and/or *Tc-kni*, we also performed simultaneous knockdown of *Tc-nub* and *Tc-cas* to determine whether they might play a redundant role in the posterior abdomen. Double *Tc-nub + Tc-cas* knockdowns do not display any posterior abdominal phenotypes, but 10/19 (52% of) cuticles examined exhibited defects in leg morphology. Specifically, the pretarsi, or claws, of the thoracic legs were almost entirely abolished (S8 Fig, A1-C). *Tc-nub* is expressed in the leg joints (S8 Fig, D), as has been observed in other insect species [25,43]. We observed that *Tc-cas* is also expressed in the developing legs, at both the proximal and distal ends (S8 Fig, D). This is, to our knowledge, the first description of *cas* function in an arthropod limb.

## Discussion

In this study, we have shown that the genes *hb*, *Kr*, *nub* and *cas* are expressed sequentially in the SAZ of *Tribolium*, as they are in *Drosophila* neuroblasts. We have also shown that *Tc*-Nub plays a role in axial patterning, acting redundantly with the abdominal gap proteins *Tc*-Gt and *Tc*-Kni to repress *Tc-Kr* expression, and thereby to establish normal abdominal Hox gene expression. Our findings provide support for the theory that the neuroblast timer network was co-opted for axial patterning.

### The neuroblast timer gene *Tc-nub* contributes to *Tc-Kr* repression in *Tribolium*

Our combinatorial knockdown experiments indicate that *Tc*-Gt, *Tc*-Kni and *Tc*-Nub all contribute to the repression of *Tc-Kr* in the abdomen. *Dm*-Gt and *Dm*-Kni are known to repress *Dm-Kr* expression in *Drosophila* [1], and *Tc*-Gt has long been suspected to regulate *Tc-Kr* expression in *Tribolium* [2,3]. However, this is, to our knowledge, the first evidence for Kni regulating *Kr* expression in *Tribolium* [35,36], and the first evidence that this role is conserved for any Kni protein outside of *Drosophila* [1]. To our knowledge this is also the first time that Nub has been shown to repress abdominal *Kr* in the context of segment patterning in any arthropod.

Our finding that *Tc*-Nub represses *Tc-Kr* expression is consistent with what is known of Nub’s function in neuroblasts in *Drosophila*. Ectopic expression of *Dm-nub* in *Drosophila* neuroblasts results in the premature termination of *Dm-Kr* expression, while *Dm-nub/Dm-pdm2* double mutants express *Dm-Kr* for an extended period of time [44,45]. Although the neuroblast timer network has not been functionally dissected in *Tribolium* neuroblasts, these cells seem to express *Tc-Kr* and *Tc-nub* sequentially [26], and it is therefore plausible that Tc-Nub also represses *Tc-Kr* expression in this context.

We propose that Nub may also regulate *Kr* expression during axial patterning in other insect species, with varying strength and/or degrees of redundancy. In *Oncopeltus*, *nub* pRNAi results in almost total loss of *abd-A* expression in the abdomen [24]. Likewise, in *Drosophila* we have observed subtle repression of *abd-A* expression in *nub/pdm2* mutants [42], in contrast to previous reports [24]. Kr is known to repress *abd-A* in *Drosophila* [46], so this effect of *nub/pdm2* on *abd-A* could be mediated by de-repression of *Kr*. If this is the case, then the interaction between Nub and Kr during segment addition may be broadly conserved across the insects.

### Redundancy in the gap gene network of *Tribolium castaneum*

*Tc-nub*, *Tc-kni* and *Tc-gt* seem to display “distributed redundancy” – that is, they have different but overlapping roles, so that if one gene is lost, the others can at least partially compensate for it (Wagner, 2005). Distributed redundancy is an efficient way to create complex, responsive gene networks that retain robustness to perturbation through mutation. There are obvious reasons why the gap gene network might benefit from being robust to mutation. These genes are responsible for regulating some of the earliest and most crucial elements of the insect body plan (segment boundaries and segment identities), and complete disruption of gap gene function is lethal [47–49]. The overlapping functions of *Tc-nub*, *Tc-gt* and *Tc-kni* may also be important for fine-tuning the expression dynamics of *Tc-Kr*, allowing for more precise regulation of the overlapping Hox gene domains in the posterior thorax and anterior abdomen.

In *Drosophila*, the posterior boundary of *Dm-Kr* is also regulated redundantly by *Dm*-Gt, *Dm*-Kni and, we suspect, *Dm*-Nub [1,42]. However, the relative contribution of these three genes to *Kr* repression varies between insect species. In *Oncopeltus*, for example, knockdown of *Oc-gt* or *Oc-kni* has no effect on *Oc-Kr* expression [50], while *Oc-nub* knockdown has severe effects on abdominal segment identity [25] and presumably on *Oc-Kr* expression (although this remains untested). In this species, then, Nub may play a more central role in the regulation of *Kr* than Gt or Kni. Conversely, in *Acheta*, *Ad-nub* knockdown has no effects on abdominal segment identity [25], suggesting that Nub has either little/no role, or a redundant role, in *Kr* repression in this species.

Subtle alterations in network interactions, even while the overall output of the network is conserved (known as developmental systems drift), are a common feature of the gap gene network [51,52], especially as adaptations to changes in upstream patterning [51,53]. Investigating the functional overlap between Nub, Gt and Kni in additional insect species, with different modes of segmentation, and at strategic points in the insect phylogeny, will help to determine when and how the function of these genes has drifted over evolutionary time. This also represents a promising framework for studying the evolutionary dynamics of gene regulatory network evolution.

Intriguingly, the phenotypes observed after simultaneous knockdown of *Tc-nub*, *Tc-kni* and *Tc-gt* are very reminiscent of those observed after knockdown of *Tc-mille-pattes* (*Tc-mlpt*). Both treatments lead to the expansion of *Tc-Kr* expression into the SAZ, and the formation of ectopic legs on presumptive abdominal segments [54]. *Tc*-Mlpt is required for expression of *Tc-gt* in the SAZ [54], so one possible explanation for this overlap in phenotypes is that *Tc*-Mlpt might also be required for expression of *Tc-kni* and *Tc-nub* in the SAZ. In this case, knockdown of *Tc-mlpt* expression would effectively phenocopy a triple knockdown of *Tc-nub*, *Tc-gt* and *Tc-kni* in the SAZ. Further investigation will be required to determine the regulatory interactions between *Tc*-Mlpt and these semi-redundant genes.

There is longstanding evidence that redundancy is common in the gap gene network of *Drosophila* [1], but our study is the first to our knowledge to explicitly test for this in the gap gene network of another species. Our findings are a reminder that failure to test for redundancy can result in network components being missed, despite their significant developmental and evolutionary relevance.

### Regulation of Hox genes in *Tribolium* – a central role for the neuroblast timer genes?

Our results suggest that *Tc*-Kr acts as a repressor of the abdominal Hox gene *Tc-Ubx*, in addition to *Tc-abd-A*. This contrasts somewhat with our current understanding of Hox gene regulation in *Drosophila* and *Tribolium*. Knockdown or mutation of *Kr* in either species leads to reduced *Ubx* expression, which has been interpreted as suggesting either that Kr acts as a direct activator for *Ubx*, or that it promotes *Ubx* expression indirectly by repressing *hb* expression [3,41,55,56]. We observe that *Tc-Ubx* expression is repressed in embryos where *Tc-Kr*, but not *Tc-hb*, is ectopically expressed in the abdomen. This finding suggests that Tc-Kr acts as a repressor, rather than an activator, for *Tc-Ubx* expression, at least at high concentrations.

The role of gap genes in regulating Hox genes is thought to predate their role in regulating the positioning of specific segment boundaries [9]. However, we lack a detailed understanding of how the gap gene network as a whole contributes to Hox gene regulation in sequentially-segmenting insects such as *Tribolium*. It is intriguing to note that the expression domains of the first three neuroblast timer genes, *Tc-hb*, *Tc-Kr* and *Tc-nub*, align approximately with the three trunk tagma in *Tribolium* (gnathum, thorax and abdomen, respectively), save that they are shifted anteriorly to align with parasegment boundaries, and *Tc-nub* covers most but not all of the abdominal parasegments (Fig 8, A1).

**Fig 8.**
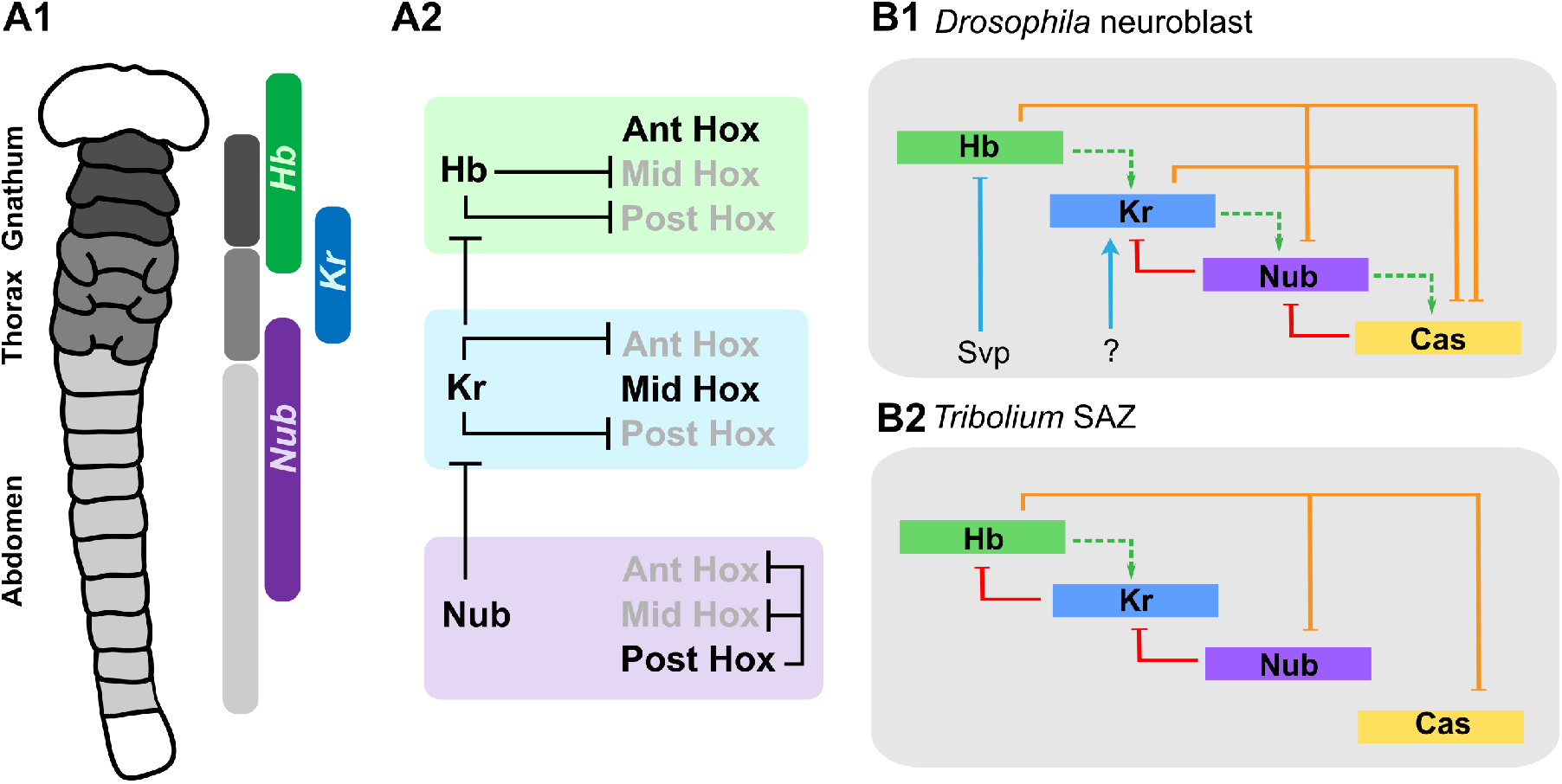
Hox and cross-regulation by the neuroblast timer proteins. **(A1)** The first three genes of the neuroblast timer series (*Tc-hb*, *Tc-Kr* and *Tc-nub*) are expressed in domains that broadly align with the three trunk tagma (the gnathum, thorax and abdomen) in *Tribolium* embryos. **(A2)** The interactions between *Tc-hb*, *Tc-Kr* and *Tc-nub* and the Hox genes are theoretically sufficient to generate three distinct domains of Hox gene expression, broadly aligning with the three major body tagma. Ant = anterior, Mid = middle, Post = posterior (referring to the position of Hox genes within the Hox cluster, and the relative position of their expression along the embryonic axis). **(B1-B2)** A summary of the interactions predicted to occur between the neuroblast timer genes in *Drosophila* neuroblasts (B1) and in the SAZ of *Tribolium* (B2). Interactions presented in B1 are based on published models [60, 61]. Svp = the nuclear transcription factor Seven-up [65]. Interactions presented in B2 are based on data from [6] and this paper. Repression of *Tc-nub* and *Tc-cas* by *Tc*-Hb has not yet been demonstrated but we infer these interactions as being likely based on their mutually exclusive expression domains in the abdomen (Fig 2). The lines representing interactions are colour-coded to represent four major ‘classes’ of interaction thought to contribute to sequential expression of the neuroblast timer genes: green = feed-forward activation; red = feedback repression; orange = “next-plus-one” repression; blue = external inputs. Note that there is some uncertainty about the relevance of “activating” interactions in this network [61], and so they have been represented with dotted lines.

Functional data also support the importance of this gene:tagma pattern (Fig 8, A2). *Tc*-Hb represses thoracic and abdominal Hox genes [6], allowing gnathal Hox genes to be expressed. *Tc*-Kr represses gnathal [3] and abdominal Hox genes, allowing the thoracic Hox genes to be expressed. Finally, *Tc*-Nub, in tandem with *Tc*-Gt and *Tc*-Kni, represses *Tc-Kr* expression, which in the absence of *Tc*-Hb allows for abdominal Hox genes to become expressed. This minimal network could therefore provide enough information to lay down the basic functional divisions of the insect axis (although not, of course, the fine details of individual segment identity).

The fact that *Tc-nub*, *Tc-gt* and *Tc-kni* could, in theory, exert their effects on Hox gene expression entirely through their interaction with *Tc-Kr* also raises the possibility that not all of the gap genes expressed during segmentation in *Tribolium* regulate Hox gene expression directly.

Taking these hypotheses further will require a molecular dissection of Hox gene regulation in *Tribolium*.

### Co-option of the neuroblast timer series for axial patterning in insects

The idea that the neuroblast timer network might be utilised for axial patterning in insects was first suggested when *Dm-hb*, *Dm-Kr*, *Dm-nub* and *Dm-cas* were found to be expressed, in that order, along the AP axis of the *Drosophila* embryo [19]. We bolster this theory by showing that the genes of the neuroblast timer network are also expressed during axial patterning in the sequentially-segmenting insect *Tribolium*, and that *Tc-nub* has a clear function during this process.

We have found that in *Tribolium*, *Tc-hb*, *Tc-Kr*, *Tc-nub* and *Tc-cas* are expressed sequentially in the cells of the SAZ, in the same order as they are in embryonic neuroblasts. Some remnant of this temporal order may also be present in *Drosophila*. Individual nuclei in the blastoderm of *Drosophila* sequentially express part of a stereotyped sequence of gap genes, with each nucleus starting from a different point in the series due to maternal pre-patterning of gap gene domains [57]. For example, many of the cells expressing *Dm-hb* will, over time, turn off *Dm-hb* expression and turn on *Dm-Kr*, while those initially expressing *Dm-Kr* will come to express *Dm-gt* [57]. These ‘damped oscillations’ cause gap gene domains to shift anteriorly across the blastoderm over time, and are thought to be a remnant of the ancestral oscillatory network that drives sequential gap gene expression in the SAZ of sequentially-segmenting insects [57]. Determining whether *Dm-nub* and *Dm-cas* are part of this sequence in *Drosophila* will require further analysis, in particular of their expression dynamics and interactions with other gap genes.

The roles of Hb, Nub and Cas in the neuroblast timer network long predate their roles in axial patterning. Homologues of all three genes (Ikaros, PouF2 and Casz1, respectively) are expressed sequentially in neural and/or retinal stem cells in mammals and promote the formation of a temporal sequence of different daughter cell types [14–17,58]. In contrast, there is no evidence that any of these genes play a role in segment formation or axial patterning outside of the arthropods. Even within the non-insect arthropods, there are species that express Hb and/or Kr in their neuroblasts but not in the SAZ [59,60]. From these observations, we can infer that at least Hb, Kr, and Nub were most likely recruited to a role in axial patterning from an ancestral role in neural patterning.

It is worth asking whether the regulatory interactions that drive sequential expression of *Tc-hb*, *Tc-Kr*, *Tc-nub* and *Tc-cas* in the SAZ are the same as in *Drosophila* neuroblasts. Sequential expression of the neuroblast timer genes in *Drosophila* neuroblasts depends largely on cross-regulatory interactions between gene products, including feed-forward activation (wherein each gene product activates the next in the series), feedback repression (wherein each gene product represses the gene that came before it) and “next-plus-one” repression (wherein each gene product represses the expression of the next-plus-one gene in the series, effectively de-repressing the next gene in the series) [12,19,61–63] (Fig 8, B1). Our understanding of the interactions between *Tc-hb*, *Tc-Kr*, *Tc-nub* and *Tc-cas* in the *Tribolium* SAZ are still fragmentary, but it does seem that this network may share some of its regulatory interactions with neuroblasts (Fig 8, B2). The time it takes for the entire sequence to be expressed in the SAZ of *Tribolium* (approximately the first half of segment addition, or 5-6 hours [64]) is also comparable to the time it takes to be expressed in neuroblasts of *Tribolium* or *Drosophila* (roughly 5-7 hours [19,26,61]).

However, there are also obvious differences between the two networks. The transition between *Tc-hb* and *Tc-Kr* expression in the SAZ appears to be mediated entirely by cross-regulatory interactions within the network [6], while in neuroblasts, the transition between *Dm-hb* and *Dm-Kr* expression is driven by an external factor, the nuclear receptor *Dm*-Seven-Up, in a cytokinesis-dependent manner [65–68]. In the SAZ of *Tribolium,* the rate of sequential activation of gap genes (including *Tc-hb* and *Tc-Kr*) is influenced by Wnt signaling, possibly mediated by the transcription factor Caudal [11] which is not expressed in the majority of neuroblasts [69]. Furthermore, the timing and extent of at least *Tc-Kr* expression in the SAZ is influenced by gap genes that are not expressed in neuroblasts, such as *Tc-gt* and *Tc-kni* [2,35; this paper]. Determining when *gt* and *kni* were incorporated into the gap gene network relative to the neuroblast timer genes will require examination of their expression and function in a broader range of insects and arthropods (taking into account that their function in axial patterning may be obscured by redundancy).

By demonstrating sequential expression of the neuroblast timer genes in the SAZ of *Tribolium*, and revealing that *Tc-nub* is able to repress the expression of *Tc-Kr* to influence Hox gene expression, our findings provide strong support for the hypothesis that the neuroblast timer network has been co-opted for axial patterning during the evolution of insects. These findings will provide a basis for future studies examining the evolution and structure of the gap gene network in insects.

## Materials and Methods

### *Tribolium castaneum* husbandry

*Tribolium castaneum* strain San Bernadino beetles (provided by A. Peel, University of Leeds, UK) were reared on organic wholemeal flour (Doves Farm Foods, Hungerford, UK) supplemented with fast action dried yeast (Sainsbury’s, London, UK) and the antifungal agent Fumagilin-B (Medivet) at 30 °C, as described in the Beetle Book v1.2 [70]. Egg lays were performed on strong white organic bread flour (Doves Farm Foods, Hungerford, UK). Incubators were maintained between 40-60 % relative humidity where possible, and no day/night cycle was used (beetles were kept in the dark).

### Collection and fixation of wild type embryos

*Tribolium* were allowed to lay on white flour for 24 hours and their eggs were then collected using a sieve with a 200 μM mesh size (Retsch test sieve 200mm x 50 mm). Collected eggs were transferred into small mesh baskets (with a mesh aperture of 250 μm) and were rinsed several times in ddH_2_O to remove all traces of flour. Their chorions were then removed by washing twice in bleach diluted with ddH_2_O to a final concentration of 2.5 % (v/v) hypochlorite, for 30-45 seconds. After further rinsing in ddH_2_O, dechorionated embryos were fixed as described by Marques-Souza et al. [6], save that a 0.68mm ID (internal diameter) needle was used to enhance devitellinisation rather than a 0.9mm ID needle. Fixed and devitellinised embryos were stored in 100 % methanol at −20 °C.

### Ovary dissection and fixation

Ovaries were removed from adult female beetles in PBS using forceps. Dissected ovaries were transferred directly into 4 % formaldehyde in PBT (PBS + 0.01 % Tween) on ice. An equal volume of heptane was added, and the tubes then rocked on a nutator for 20 minutes to allow for fixation. The ovaries were then rinsed several times in PBT and then washed into 100% methanol for storage at −20 °C.

### RNA interference

Plasmids containing clones for *GFP*, *Tc-nub*, *Tc-cas, Tc-gt* and *Tc-kni* were provided by A. Peel and R. Sharma. dsRNA was synthesised from PCR products using T7 polymerase, and was purified using phenol chloroform precipitation. Purified dsRNA was resuspended in RNase-free water and injected into *Tribolium* adults or eggs at a concentration of 1-4 μg/μL. Unless specified otherwise, single knockdowns were carried out using 2 μg/μL of dsRNA, while double and triple knockdowns used the component dsRNAs mixed to a final concentration of 1 μg/μL each (the viscosity of the injection fluid became difficult to work with above 4 μg/μL).

All injections for RNAi were carried out using a Pico-injector system (Medical Systems Corporation). Parental RNAi was carried out by injecting dsRNA into the dorsal surface of the abdomen under the elytra of adult female beetles as described by Posnien et al [71]. Males were introduced to the injected females the day after injection, and eggs were collected starting from one week after injection. Eggs were collected and fixed regularly (every 1-2 days) as described above for 3-4 weeks after injection.

Embryonic microinjection for eRNAi was carried out using a method adapted from Benton [29]. 1-2 hour old eggs were transferred into small mesh baskets (with a mesh aperture of 250 μm) and rinsed several times in ddH_2_O. Chorions were removed by washing twice in bleach, diluted with ddH_2_O to a final concentration of ~0.6% (v/v) hypochlorite, for 30-45 seconds. Eggs were rinsed again and then healthy-looking eggs were lined up on coverslips and allowed to dry. Eggs were injected into the anterior pole (to reduce the risk of damage posterior segment addition zone) in a 1:1 mix of Halocarbon oil 700 and Halocarbon oil 27 (Sigma Aldrich). The coverslip was turned over on to a Lumox culture dish as described by Benton [29], save that glass ‘feet’ approximately 0.6mm high (made from strips of #1.5 coverslip) were attached to the coverslip at either end of the injected rows of eggs, to prevent them from being pressed against the membrane. Injected eggs were then stored in plastic chambers with wet paper towel (to maintain humidity) and reared at 30 °C.

For fixation, injected embryos were aged for the appropriate length of time then injected with PBT + 10 % formaldehyde (v/v) and left to fix at room temperature for one hour. They were then transferred using an eyelash hair to Eppendorf tubes and fixed for an additional hour in a 1:1 mix of heptane and PBT + 4 % formaldehyde (v/v). The aqueous layer was removed, and 100 % ice-cold methanol added. Germbands were manually dissected away from the remainder of the yolk, chorion and vitelline membrane in PBS, and then stored in 100 % methanol at −20 °C until required.

### Hybridisation Chain Reaction (HCR) *in situ* hybridisation (ISH)

Version 3.0 HCR probes (20 pairs per gene) and fluorescently-labelled hairpins were produced by Molecular Instruments. Probe template sequences were taken from NCBI (*Tc-hb*: NM_001044628.1, *Tc-Kr*: NM_001039438.2, *Tc-nub*: XM_015979462.1, *Tc-cas*: XM_015980923.1, *Tc-wg*: NM_001114350.1, *Tc-Ubx*: XM_008203013.2, *Tc-abd-A*: NM_001039429.1). All required buffers were made according to the instructions provided by Molecular Instruments, with the one exception that the percentage of dextran sulfate in the probe hybridisation and amplification buffers was reduced from 10 % (v/v) to 5 % (v/v) to reduce viscosity and improve retention of embryos during washes.

Fixed embryos or ovaries were prepared for HCR ISH by removing methanol and replacing it with 1 mL of PBT containing 4 % formaldehyde. Tubes were rocked on the nutator for 30 minutes to allow for additional fixing and rehydration to occur. The HCR ISH was then carried out as per the Molecular Instruments HCR v3.0 protocol for whole-mount fruit fly embryos, with the exception that hybridisation steps were carried out in 100 rather than 200 μL of hybridisation buffer, and the volume of probe added was adjusted to give the same final concentration (4 nM/mL). Additionally, 1 ng/μL DAPI was added to the first 30 minute wash on the final day so that nuclear staining could be carried out in parallel. After washing, embryos or ovaries were transferred first into 25 % (v/v) glycerol and then into 50 % (v/v) glycerol before being stored at 4 °C to stiffen and clear for mounting.

### Mounting and imaging of embryos

Blastoderm stage embryos were mounted in glass-bottomed petri dishes (Cellvis), and dissected germbands and whole ovarioles on glass slides, in ProLong™ Gold Antifade Mountant (Thermofisher Scientific) as per the manufacturer’s instructions. Mounted embryos were imaged using an Olympus FV3000 confocal microscope at the Department of Zoology Imaging Facility (University of Cambridge).

### Preparation and imaging of cuticles

Embryos and larvae were processed for cuticle preparation either upon hatching, or after 7-10 days if they failed to hatch in this time. Embryos and larvae of uninjected embryos were first rinsed in 2.5 % (v/v) bleach and then in ddH_2_O to remove any remaining chorion and debris. Injected embryos and larvae were dissected out of their chorions manually, and washed in methanol and then heptane (one hour each) to remove the halocarbon oil. Embryos or larvae were then transferred to a glass slide, covered with a 1:1 mix of Hoyer’s medium [72]:lactic acid and a coverslip, and heated at 60 °C overnight. Cuticles were imaged with an excitation wavelength of 633 nm on an Olympus FV3000 in the Department of Zoology (University of Cambridge).

### Image processing and figure assembly

Images and Z stacks were stitched using the Olympus FV3000 software. Additional image processing was carried out in Fiji [73]. Fiji was used to adjust image brightness and contrast, and to rotate, crop and reslice images where necessary. Figures were assembled in the open source vector graphics editor Inkscape (https://inkscape.org/).

## Acknowledgements

We thank A. Peel and E. Clark for discussions that helped to inspire this project, and for advice and input throughout the work. We also thank A. Peel and R. Sharma for providing beetles and plasmids and for help with rearing and knockdowns. Finally, we thank S. Taylor for assistance with ovary dissection.

## Notes

### Competing Interest Statement

The authors have declared no competing interest.

