## Supplementary Material for "The neuroblast timer gene *nubbin* exhibits functional redundancy with gap genes to regulate segment identity in *Tribolium*"

### Supplementary information

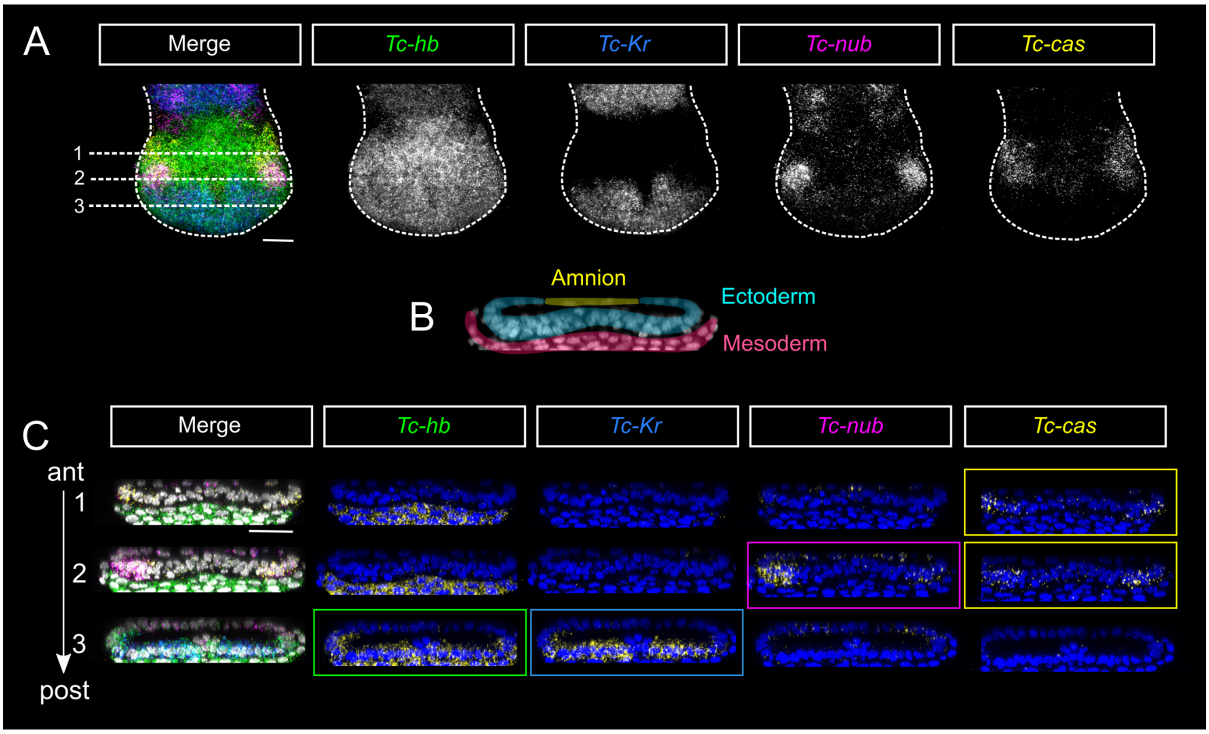


**S1 Fig. *Tc-hb*, *Tc-Kr*, *Tc-nub* and *Tc-cas* are all expressed in the ectoderm at the posterior-most end of the embryo (the presumptive hindgut). A)** Maximum projection of confocal z-stacks spanning the posterior SAZ after segment addition is completed, but before gut morphogenesis begins. *Tc-hb*, *Tc-Kr*, *Tc-nub* and *Tc-cas* are expressed in overlapping domains in this region. Anterior is to the top, and ventral is along the vertical midline. **B)** A transverse section of the posterior SAZ showing the arrangement of amnion, ectoderm and mesoderm (as judged by tissue morphology) in false colours. Dorsal is to the top. **C)** Transverse sections of the posterior SAZ from the same embryo shown in A) at three positions along the anterior-posterior axis (labelled as 1, 2 and 3). At position 1, the most anterior position, only *Tc-cas* is expressed in the ectoderm. At position 2, the central position, both *Tc-nub* and *Tc-cas* are expressed in the ectoderm. Finally, at position 3, the most posterior position, *Tc-hb* and *Tc-Kr* are expressed in the ectoderm. *Tc-hb* was expressed in the mesoderm throughout the SAZ. Dorsal is to the top. Each of the maximum projections in panels B and C spans approximately 5-10 µM along the anterior-to-posterior axis of the embryo. Scale bar = 20 µM.

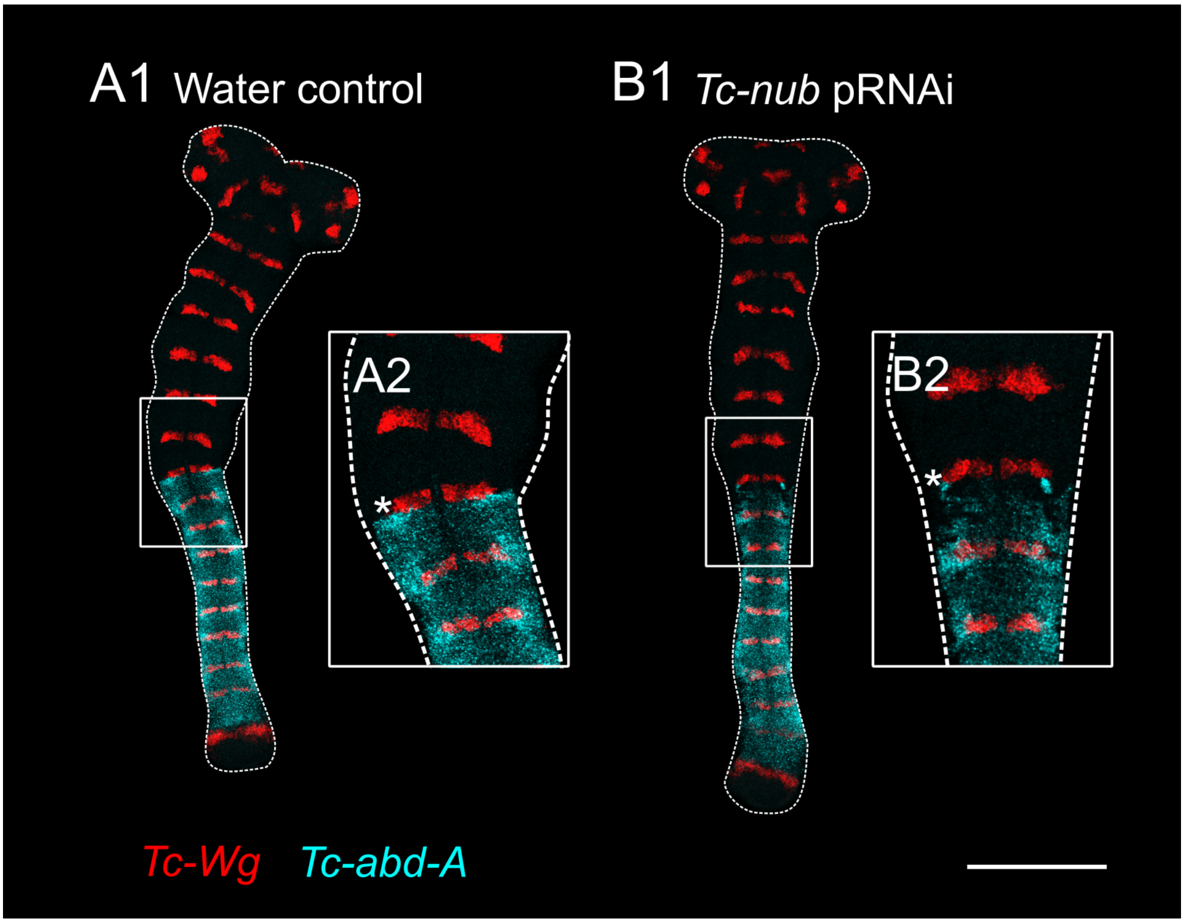


**S2 Fig. Expression of the Hox gene *Tc-abd-A* in parasegment 7 is disrupted following *Tc-nub* pRNAi.** **(A1-A2)** Embryos produced by mothers injected with water showed normal expression of *Tc-abd-A* in the anterior abdomen, abutting the posterior of wg6 (marked by an asterisk). **(B1-B2)** Embryos produced by mothers injected with *Tc-nub* dsRNA (2 µg/µL) showed disrupted *Tc-abd-A* expression in the anterior of parasegment 7, just posterior to wg6 (marked with an asterisk)**.** Both images are maximum projections of confocal z-stacks through dissected, flat mounted embryos. Anterior is to the top and ventral is along the vertical midline of the embryo. Scale bar = 200 µM.

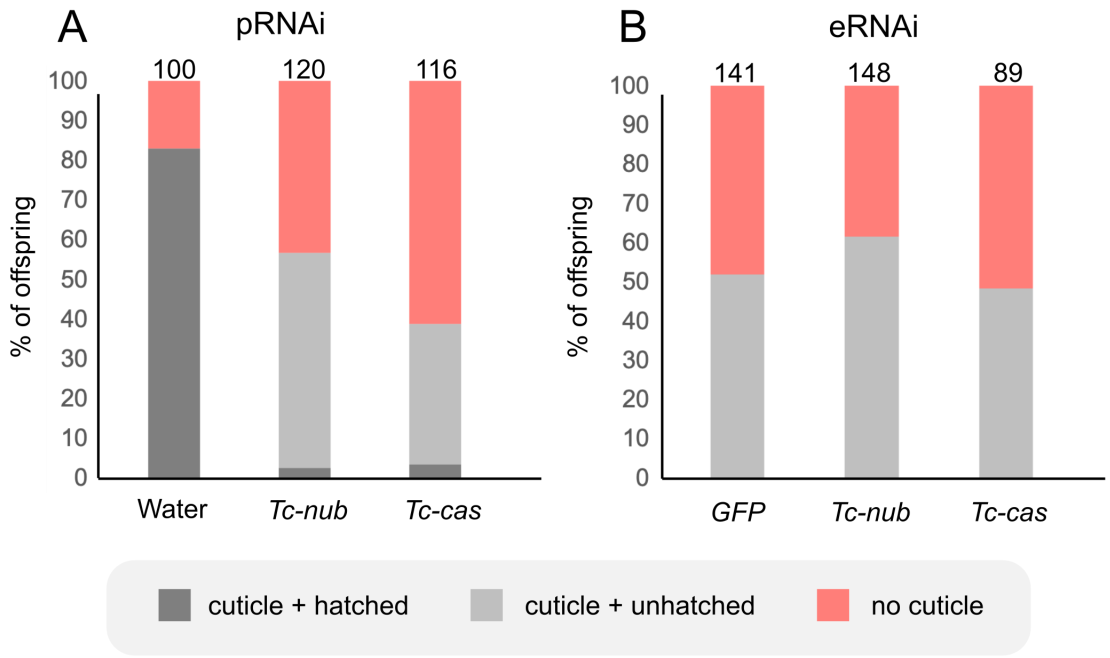


**S3 Fig. Percentage of embryos hatching and developing to the point of cuticle formation in control, *Tc-nub* and *Tc-cas* RNAi experiments. (A)** pRNAi against *Tc-nub* or *Tc-cas* (2 µg/µL) reduced the percentage of eggs forming cuticle from ~80% (in water-injected controls) to less than 50%. Furthermore, many of the eggs that did form apparently normal cuticle after *Tc-nub* or *Tc-cas* pRNAi failed to hatch. Note that water and *GFP* controls gave similar results for pRNAi (Table S4). **(B)** After eRNAi, the percentage of eggs forming cuticle was similar in *GFP* controls compared to *Tc-nub* or *Tc-cas* knockdowns (all dsRNAs injected at 2 µg/µL). Hatching rates were not recorded for eRNAi as maintenance of embryos in halocarbon oil supressed hatching in all treatments.

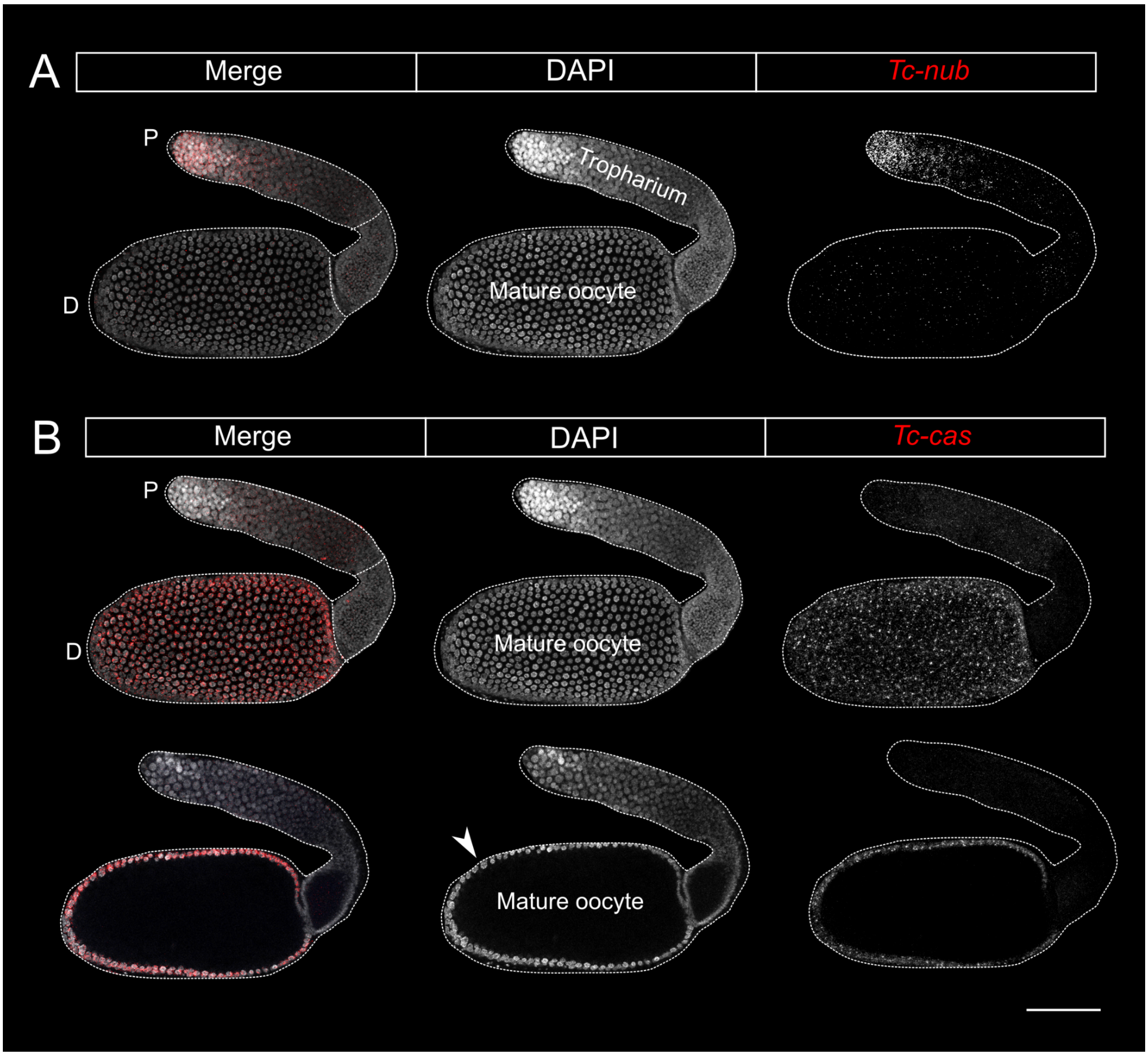


**S4 Fig. Expression of *Tc-nub* and *Tc-cas* in the ovarioles of mature female *Tribolium*. (A)** *Tc-nub* is expressed in a subset of nurse cells in the proximal tropharium. **(B)** *Tc-cas* is expressed in the follicular cells that surround the mature oocyte. The upper row of images show a maximum projection through an entire dissected ovariole, while the lower row of images show a maximum projection through ~10 µM of the ovariole's center, illustrating that *Tc-cas* expression is limited to the outer layer of follicular cells (white arrowhead) and is absent in the oocyte itself. Panels are from the same image, and are maximum projections of z-stacks through a dissected ovariole. P = proximal end of the ovariole, D = distal end of the ovariole. Scale bar = 100 µM.

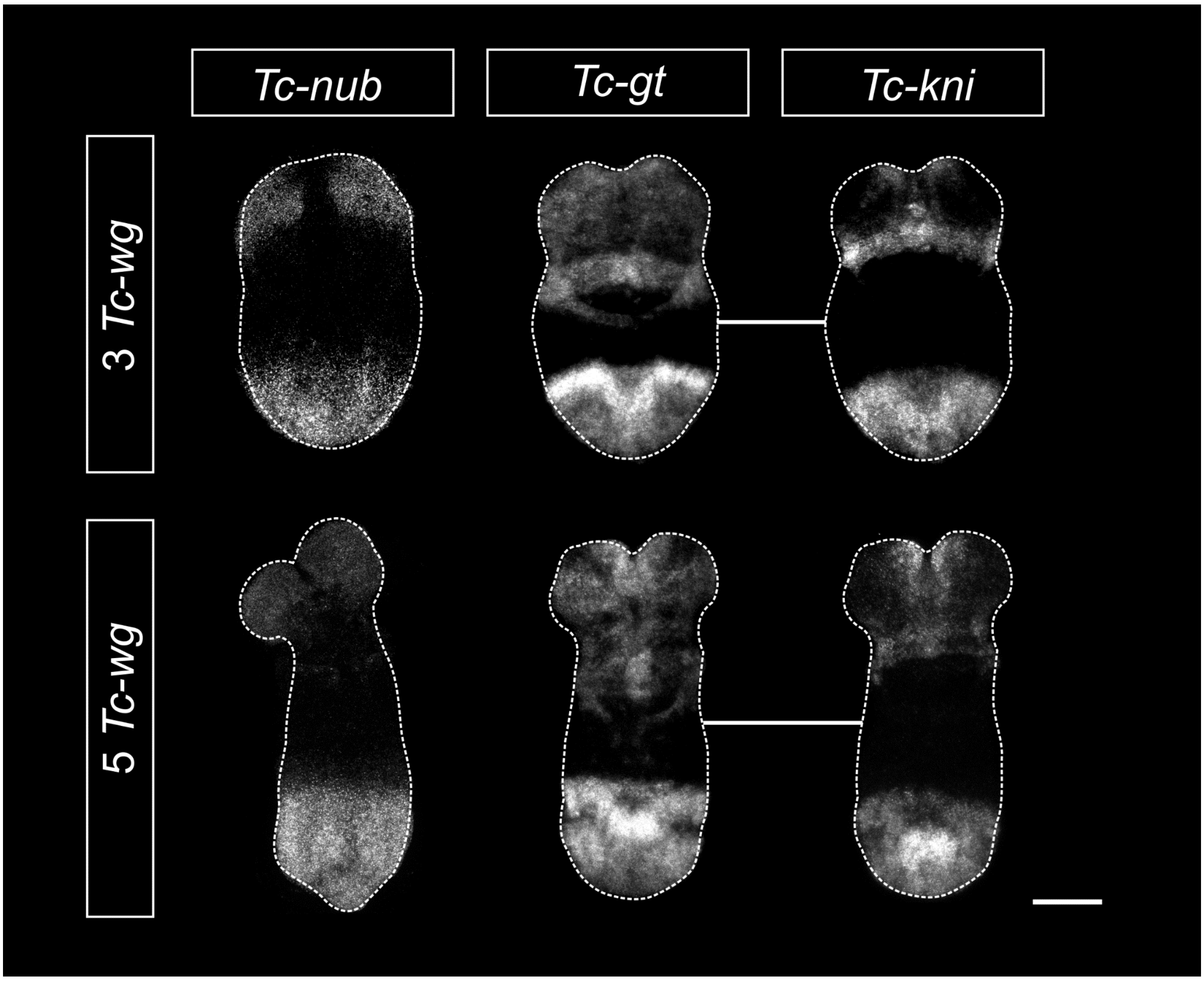


**S5 Fig. Overlapping expression domains of *Tc-nub*, *Tc*-gt and *Tc-kni* in the SAZ of embryos with three trunk *Tc-wg* stripes and five trunk *Tc-wg* stripes.** At both stages, *Tc-gt* and *Tc-kni* images were taken from the same embryo, indicated by a white line joining them. Images are maximum projections through flat mounted, dissected germbands. Scale bar = 100 μM.

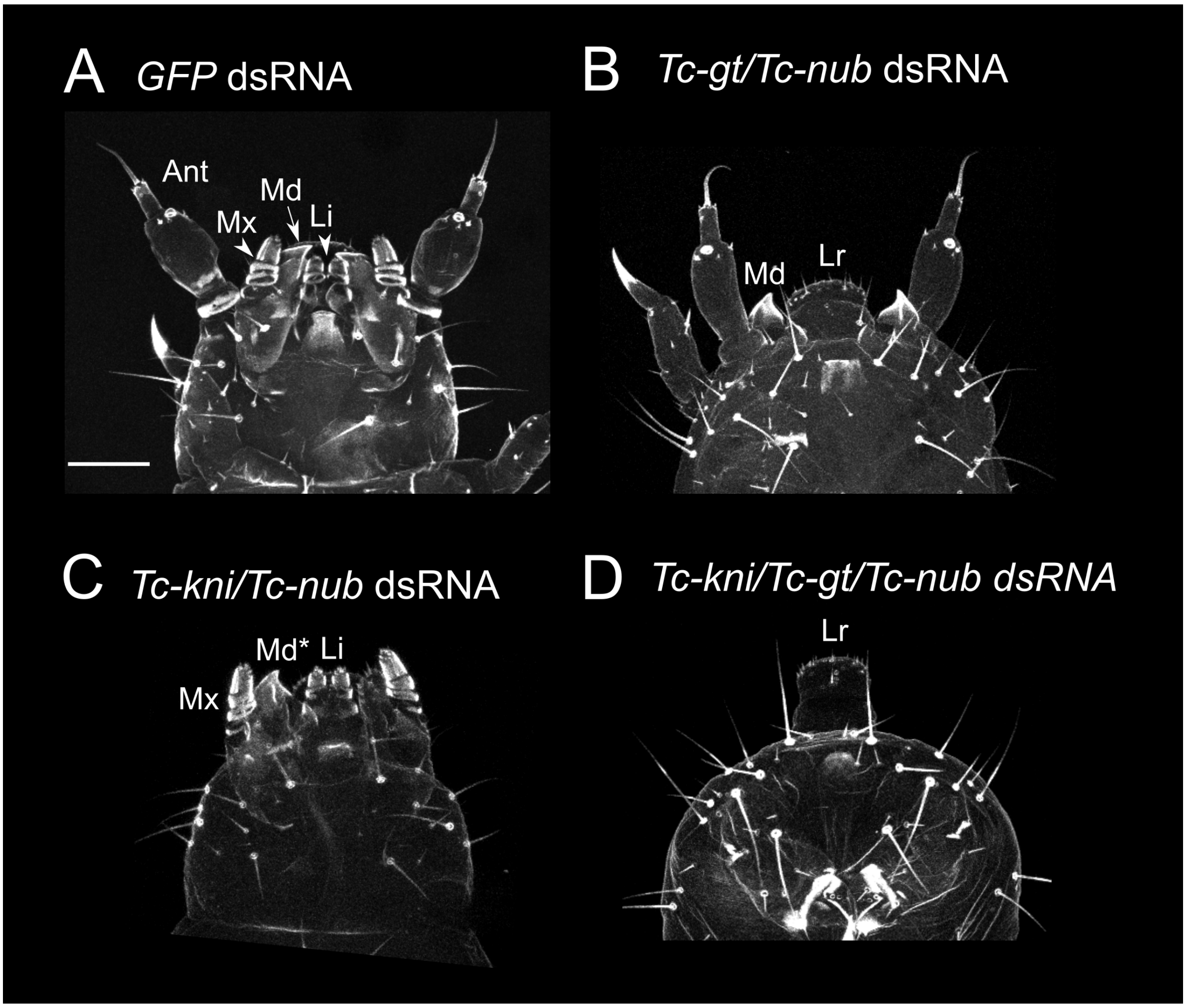


**S6 Fig. *Tc-nub* eRNAi did not enhance the effects of *Tc-gt* or *Tc-kni* knockdown on external head development. (A)** Embryos injected with *GFP* dsRNA (2 µg/µL) had wild type external head morphology. **(B)** In embryos injected with *Tc-nub* + *Tc-gt* dsRNA (1 µg/µL each), the maxillae and labium were transformed into legs, while the mandibles, antennae and labrum were left intact, as observed in *Tc-gt* single knockdowns. **(C)** In embryos injected with *Tc-kni* + *Tc-nub* dsRNA (1 µg/µL each), the antennae and one or more mandibles was lost, but the maxillae, labium and labrum remained intact, as observed in *Tc-kni* single knockdowns. (D) Embryos injected with *Tc-kni* + *Tc-gt* + *Tc-nub* dsRNA (1 µg/µL each) displayed an additive phenotype; the antennae and mandibles are lost, while the maxillae and labium are transformed into legs. These data suggest that *Tc-nub* does not act redundantly with *Tc-kni* and/or *Tc-gt* to regulate head development in *Tribolium*. An = antenna; Md = mandible; Mx = maxilla; Li = labium; Lr = labrum. In C, Md* indicates the single remaining mandible (the second mandible is lost in this knockdown). All images are maximum projections of confocal z-stacks through cuticle preparations. Scale bar is 50 µM.

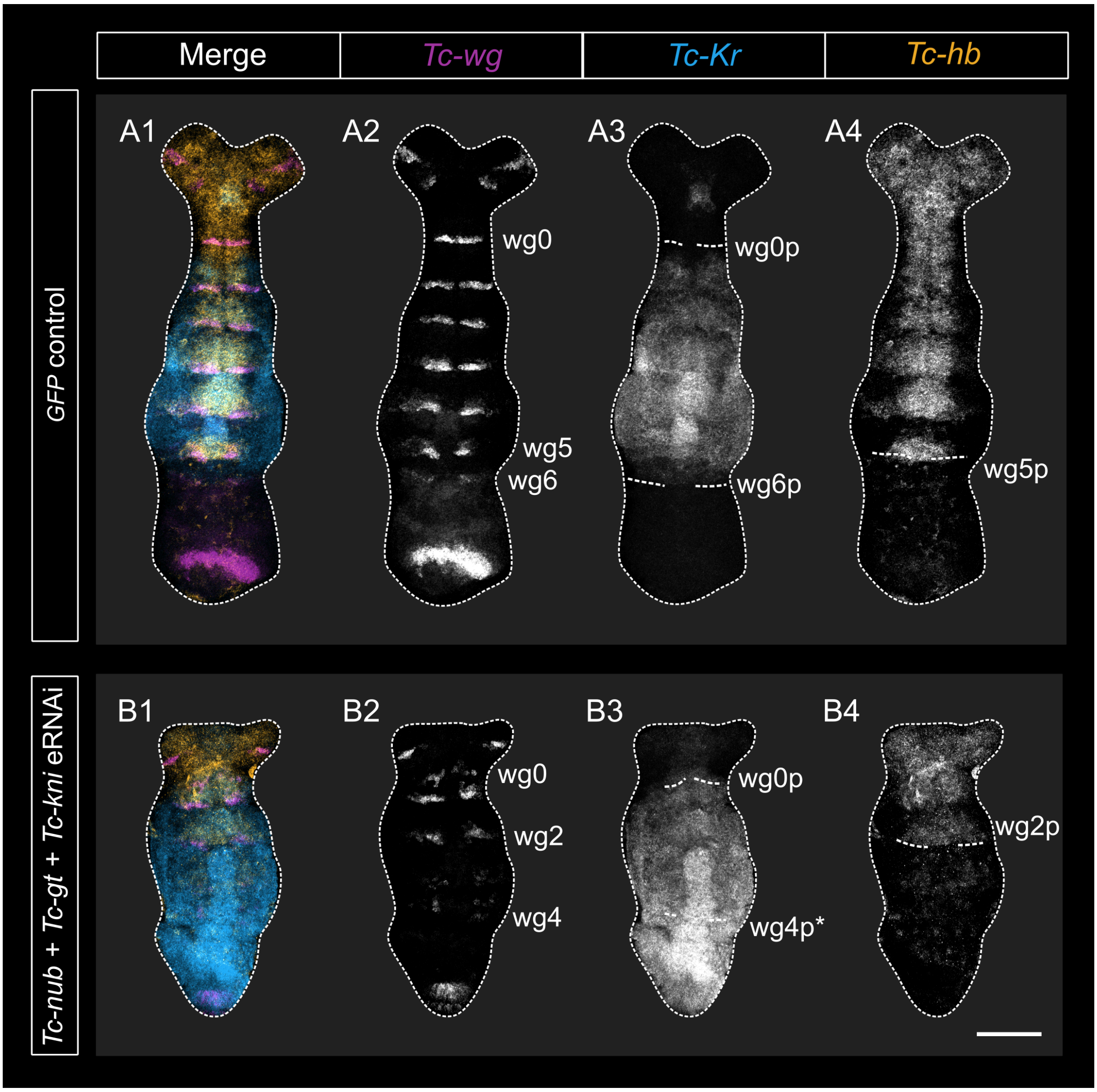


**S7 Fig. Expression of *Tc-Kr,* but not *Tc-hb,* was expanded posteriorly after eRNAi against *Tc-nub* + *Tc-gt* + *Tc-kni.*** **(A1-A4)** In *GFP* dsRNA-injected control embryos the expression of *Tc-hb* and *Tc-Kr* matched descriptions of wild-type expression [3,28]. **(B1-B4)** In embryos injected with *Tc-nub* + *Tc-gt* + *Tc-kni* dsRNA, *Tc-Kr*, but not *Tc-hb,* expression was expanded compared to similarly staged wild type embryos [3,28]. Embryos were fixed 16-17h AEL. All embryos were imaged using the same laser settings and brightness/contrast values were adjusted identically for all images. All images are maximum projections through flat mounted, dissected germbands. Anterior is to the top and ventral along the vertical midline of each embryo. wg0-6 = *Tc-wg* stripes 0-6; wg0-6p = posterior boundary of *Tc-wg* stripes 0-6. Scale bar is 100 µM.

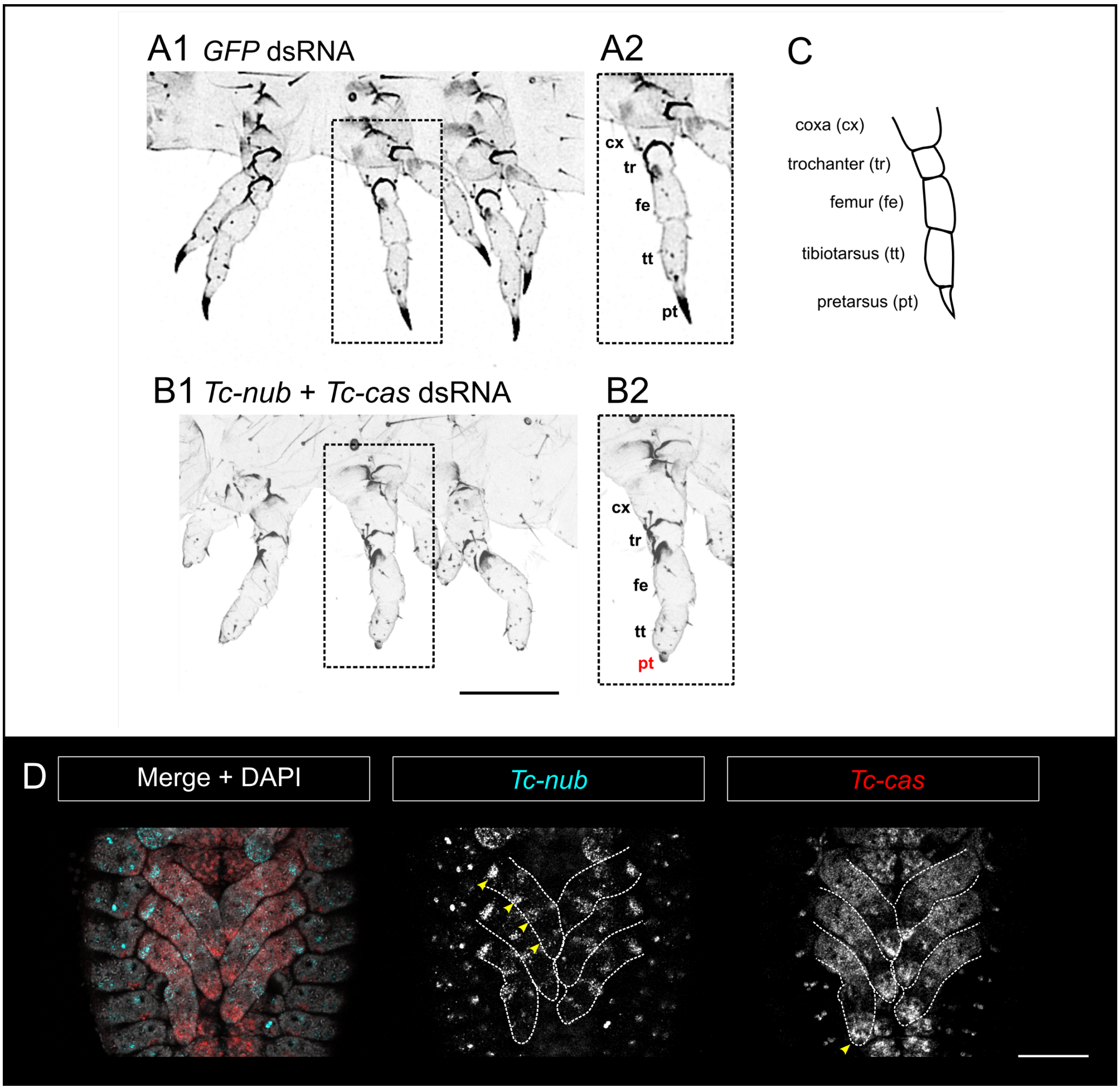


**S8 Fig. *Tc-nub* and *Tc-cas* are required for pretarsus development. (A1-A2)** Embryos injected with *GFP* dsRNA (2 µg/µL) had wildtype external leg morphology. **(B1-B2)** In embryos injected with *Tc-nub* + *Tc-cas* dsRNA (1 µg/µL each), the most distal leg segment - the pretarsus (pt) – failed to form normally. **(C)** A graphical summary of the leg segments in *Tribolium*. **(D)** Expression of *Tc-nub* and *Tc-cas* in the developing legs. Yellow arrowheads mark the rings of *Tc-nub* expression in the presumptive leg joints, and the expression of *Tc-cas* in the most distal portion of the leg, where the pretarsus will form. Images in A1-B2 and D are maximum projections of confocal z-stacks through cuticle preparations and dissected, flat mounted germbands, respectively. Scale bars are 100 µM.

**S1 Table. Cuticle phenotypes following pRNAi against *GFP*, *Tc-odd*, *Tc-nub* or *Tc-cas*.** *Tc-odd* dsRNA was used as a positive control, and the phenotypes observed in offspring were consistent with those observed in previous publications (Choe et al., 2006). At 2 μg/μL, *Tc-nub* knockdown produced a range of cuticle phenotypes at low frequency, mostly affecting segment formation and patterning in the abdomen. Only the 'nub' phenotype was investigated in detail in this paper, as other phenotypes were not consistently identified in eRNAi experiments. The small percentage of cuticle defects observed after *Tc-cas* pRNAi at 2 μg/μL were not consistent between experiments and were not investigated further. N = number of eggs examined; WT = wild type; # (%) nubs = number (and percentage in brackets) of embryos that developed ectopic cuticular protrusions on abdominal segment 1.

| **dsRNA injected** | **N** | **# (%) hatching** | **# (%) WT cuticles** | **# (%) nubs** | **# (%) other cuticle defects** | **# (%) no cuticle** |
|---|---|---|---|---|---|---|
| *GFP* (1 μg/μL) | 404 | 317 (78) | 336 (83) | 0 (0) | 0 (0) | 68 (17) |
| Water | 100 | 82 (82) | 83 (83) | 0 (0) | 0 (0) | 17 (17) |
| *Tc-odd* (1 μg/μL) | 116 | 0 (0) | 0 (0) | 0 (0) | 49 **(42)** | 67 **(58)** |
| *Tc-nub* (1 μg/μL) | 447 | 16 (4) | 198 (44) | 0 (0) | 0 (0) | 249 **(56)** |
| *Tc-nub* (2 μg/μL) | 120 | 3 (3) | 60 (50) | 2 **(2)** | 6 (5) | 52 **(43)** |
| *Tc-cas* (1 μg/μL) | 167 | 8 (5) | 68 (41) | 0 (0) | 0 (0) | 99 **(59)** |
| *Tc-cas* (2 μg/μL) | 116 | 4 (3) | 44 (38) | 0 (0) | 1 (<1) | 71 **(61)** |

**S2 Table. Cuticle phenotypes following eRNAi against one or more of the genes *Tc-nub*, *Tc-kni* and *Tc-gt*.** Single knockdowns were carried out using 2 µg/µL of dsRNA, while all double and triple knockdowns used the component dsRNAs mixed to a final concentration of 1 µg/µL each. N = number of eggs injected; '# (%) cuticles' = the number (and percentage, in brackets) of injected embryos going on to form cuticles; '# (%) Abdominal transformations' refers to the number (and percentage, in brackets) of cuticles in which abdominal segments have been transformed towards a thoracic fate; 'nubs' and 'legs' refer to cuticular protrusions without joints/claws, or partial/complete legs (with joints and/or claws), respectively; 'Avg (max) No. extra leg pairs' refers to the average (and maximum) number of pairs of legs presumed to have formed through homeotic transformation of an abdominal segment.

| Ttreatment (dsRNA injected) | | N | # (%) cuticles | # (%) Abdominal transformations | | | Avg (max) No. extra leg pairs |
|---|---|---|---|---|---|---|---|
| | | | | 'nubs' | legs | Total | |
| **Singles** | *GFP* | 266 | 171 (64) | 0 (0) | 0 (0) | 0 (0) | 0 (0) |
| | *Tc-nub* | 148 | 91 (62) | 11 (12) | 0 (0) | 11 **(12)** | 0 (0) |
| | *Tc-kni* | 45 | 28 (62) | 0 (0) | 0 (0) | 0 (0) | 0 (0) |
| | *Tc-gt* | 50 | 36 (72) | 0 (0) | 4 (11) | 4 **(11)** | **1 (1)** |
| **Doubles** | *Tc-nub* + *Tc-kni* | 93 | 41 (44) | 18 (43) | 10 (24) | 28 **(68)** | **1 (1)** |
| | *Tc-gt* + *Tc-kni* | 49 | 28 (57) | 7 (25) | 13 (46) | 20 **(71)** | **1 (1)** |
| | *Tc-nub* + *Tc-gt* | 95 | 38 (46) | 12 (31.6) | 19 (50) | 31 (**82)** | **1.3 (2)** |
| **Triple** | *Tc-nub* + *Tc-gt* + *Tc-kni* | 136 | 35 **(26)** | 0 (0) | 0 (0) | 33 **(94.3)** | **4.0 (7)** |
